# Whole-Brain Co-Mapping of Gene Expression and Neuronal Activity at Cellular Resolution in Behaving Zebrafish

**DOI:** 10.64898/2026.02.07.704095

**Authors:** Emmanuel Marquez-Legorreta, Greg M. Fleishman, Luuk W. Hesselink, Mark Eddison, Kasper Smeets, Carsen Stringer, Philipp J. Keller, Sujatha Narayan, Alex B. Chen, Brett D. Mensh, Scott M. Sternson, Bernhard Englitz, Paul W. Tillberg, Misha B. Ahrens

## Abstract

The brain’s capabilities rely on both the molecular properties of individual cells and their interactions across brain-wide networks. However, relating gene expression to activity in individual neurons across the entire brain remains elusive. Here we developed an experimental-computational platform, WARP, for whole-brain imaging of neuronal activity during behavior, expansion-assisted spatial transcriptomics, and cellular-level registration of these two modalities. Through joint analysis of whole-brain neuronal activity during multiple behaviors, cellular gene expression, and anatomy, we identified functions of molecularly defined populations—including luminance coding in a *cckb-pou4f2* midbrain population and task-structured activity in *pvalb7-eomesa* hippocampal-like neurons—and defined over 2,000 other function-gene-anatomy subpopulations. Analysis of this unprecedented multimodal dataset also revealed that most gene-matched neurons showed stronger activity correlations, highlighting a brain-wide role for gene expression in functional organization. WARP establishes a foundational platform and open-access dataset for cross-experiment discovery, high-throughput function-to-gene mapping, unification of cell biology and systems neuroscience, and scalable circuit modeling at the whole-brain scale.

## Introduction

The computations that give rise to perception, memory, and behavior emerge from interactions among diverse cells across the entire brain.^1^ The cellular processes underlying these interactions are shaped by the genes expressed in individual cells. A frontier in neuroscience is to unify whole network dynamics with the cell-biological understanding of neurons by capturing the molecular identity, cell state, and activity of every cell across the entire central nervous system, during behavior, in the same animal. As demonstrated by genetic approaches in highly tractable model organisms, cell-type specific marker genes facilitate addressing mechanistic circuit questions and replication across animals.^2–4^ Integrating multi-gene expression profiles with time-varying activity and anatomy^5^ across the complete brain—thus creating a joint definition of functional-genetic-anatomical cell type—would reveal how brain function and animal behavior arise from the molecular makeup of all constituent individual cells.

While previous studies have linked neuronal dynamics to gene expression in localized brain regions,^6–9,5,10^ methods operating at the whole-brain scale for both activity and cell molecular identity in individual animals are lacking. This is due to the challenges in recording both activity and gene expression modalities across the entire brain and then registering them both at cellular resolution. However, recent advancements in multiplexed RNA transcript visualization have enabled high-dimensional gene expression profiling at the single-cell level across large tissue samples,^11–15^ including further development of expansion microscopy methods that facilitate sub-micron scale imaging in large 3D volumes.^12^ In this context, larval zebrafish provide a unique opportunity to apply these developments at the whole-brain scale due to their accessible size and the ability to record the activity of nearly all neurons alongside sensory stimulation and behavior.^16,17^ That said, integrating time-varying activity during behavior with RNA measurements across the entire brain of an animal presents a significant challenge.

The integration of gene expression, neuronal dynamics, and anatomy requires bridging the fine spatial scale necessary for sub-micron RNA transcript assignment, to cellular activity measurements, to whole-brain dimensions, at single-cell precision. The scale of this task includes aligning terabyte-sized transcriptomic images with whole-brain functional data, alongside behavioral analysis, signal detection of neuronal activity, accurate transcript assignment and multimodal volumetric image registration. Importantly, these methods should maintain a high reliability across the whole brain volume of over a hundred thousand tightly packed neurons while keeping the computational burden bounded.

Here, we provide a complete framework of experimental and computational methods that, for the first time, enables the acquisition and alignment of functional, transcriptomic, and anatomical data at single-cell accuracy of most cells across the whole brain. This integrated suite of methods, termed WARP (Whole-brain neuronal Activity and RNA Profiling), combines light-sheet microscopy to record calcium activity during behavioral assays, expansion microscopy to measure the expression of dozens of genes in the same brain with high spatial resolution and optimized RNA probe accessibility. We further developed a suite of multiple novel open-source computational methods to register and extract function-gene-anatomy information from the acquired data. In addition to whole-brain coverage, this endeavor resulted in a combined function-gene-anatomy throughput that was 200-300x higher than previous studies, with up to 77,000 neurons per brain containing both functional and gene expression profiles.

Our method generated a unique multimodal open-access dataset with hundreds of thousands of cells across a main set of 3 animals and a secondary set of 3 animals, where we classified hundreds of genetically defined subtypes of neurons throughout the brain, each with associated activity traces during behavior, and over 2,000 functional-genetic-anatomically defined subpopulations.

This work will impact neuroscience research in several important ways. It shifts the focus from traditional region-by-region studies to a broader and more exhaustive understanding of the full plethora of molecular cell types relevant to specific behaviors at the level of single cells, across whole brains, in individual animals. It improves replicability in systems neuroscience by aligning cell types across individuals and enabling high-throughput gene screening to choose transgenic lines to create for targeted perturbation studies. Overall, it will help us better understand how behavior, activity, molecular identity and anatomy are linked throughout the entire brain.

## Results

### Combined calcium imaging, expansion microscopy, and multiplexed *in situ* hybridization through experimental and computational advances

To obtain a dataset that combines activity and gene expression^5,18^ at the whole-brain, single-cell level, we first recorded neuronal activity in larval zebrafish while they engaged in visual behavioral assays. We presented multiple assays as sessions during one experiment to maximize the range of brain circuits engaged during the imaging session. Assays included the optomotor response (OMR) encouraging fish to swim along with visual motion in different directions and at varying speeds,^19,20^ stimuli with changes in luminance^21^ and simulated looming predators,^22–24^ a futility-induced passivity paradigm,^25^ as well as other behaviors evoked by visual stimuli. Swimming was recorded directly from spinal motor nerves using non-invasive electrodes touching the skin in a system previously developed for virtual-reality behavior.^16^ Neuronal activity was detected by imaging nuclear-targeted GCaMP7f^26^ in almost all neurons in the brain of *Tg(elavl3:H2B-GCaMP7f)* transgenic fish using whole-brain light-sheet microscopy^16^ (Figure 1A,B, Video S1) (see Methods: Fictive behavior; Calcium imaging).

**Figure 1.**
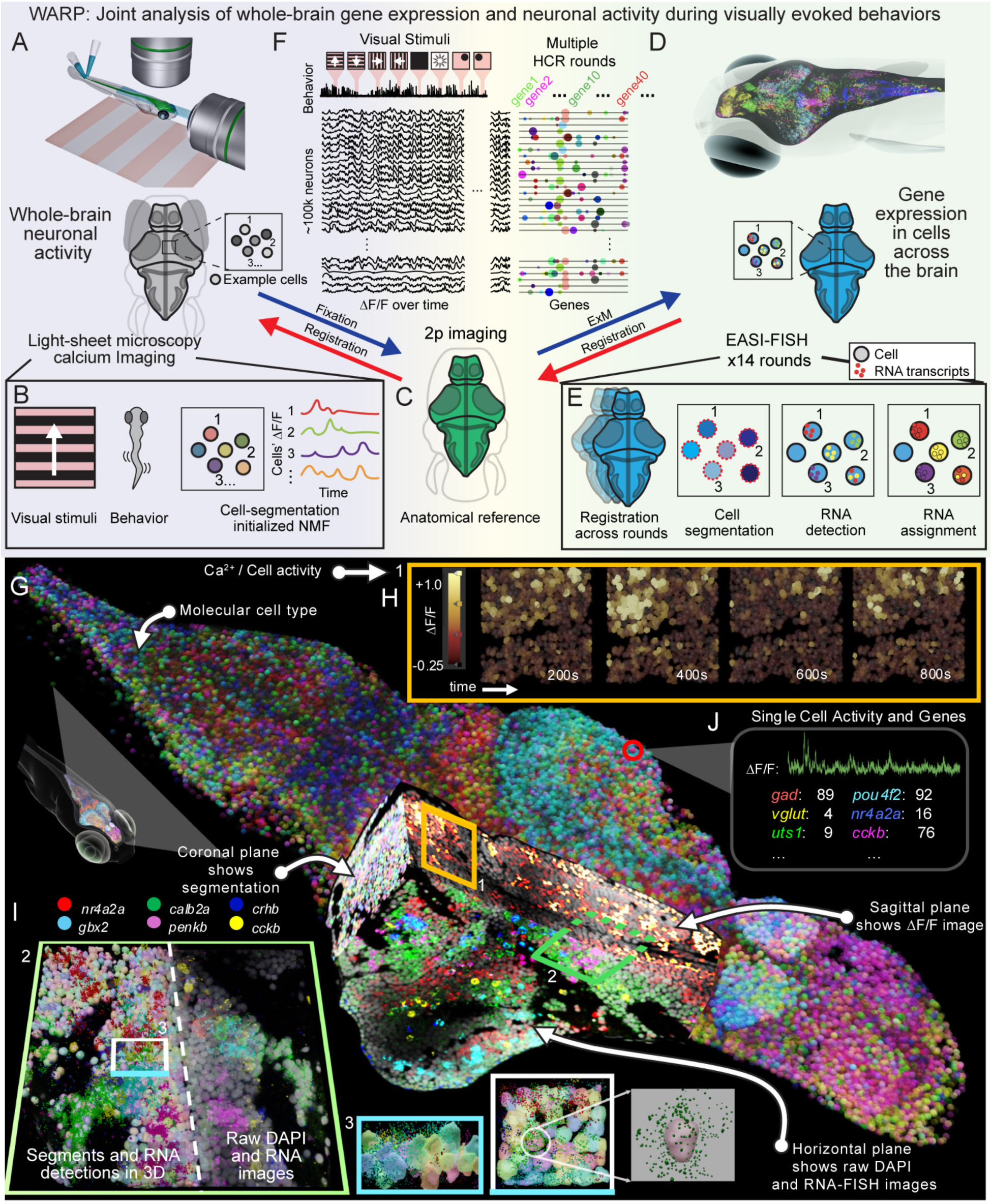
The WARP platform correlates behavior to whole brain calcium imaging and spatial transcriptomics, with single-cell accurate registration, segmentation and RNA assignment, in the same animal. (A-F) WARP workflow using experimental procedures, and algorithms presented in Figure 2. (A) Fish were presented with diverse visual stimuli during whole brain calcium imaging and neuromuscular electrophysiology to register fictive behavior. (B) Calcium imaging data was processed with a novel segmentation-initialized non-negative matrix factorization (NMF) resulting in an individual activity trace per neuron. (C) After functional imaging the sample was fixed, and an anatomical reference image was acquired using two-photon microscopy. GCaMP and/or jRGECO expression was used to register across modalities. (D) The gelled fish brain then underwent multiple rounds of EASI-FISH to detect RNA transcripts. (E) The multiple EASI-FISH rounds were then registered to each other. Cells were segmented using DAPI nuclear stain. Individual RNA transcripts were then detected and assigned to a cell segment using a novel RNA spot assignment algorithm. (F) Illustration of the resulting multimodal dataset that includes behavioral responses, activity traces of thousands of neurons, and mRNA transcript levels of these neurons. (G) A 3D illustration of a fully processed sample. Extracted cells yield a neural activity trace and transcript count for 41 genes. Cells are colored based on their gene expression combinations. The 3D model cutaways show raw RNA transcript images (horizontal), standard deviation of neural activity over 20 min (sagittal), and segmented cells (coronal). (H) Changes in calcium fluorescence in a crop of the brain. (I) A horizontal top view of the thalamus, showing segmented cells and detected RNA spots (left), and the DAPI stain with multiple rounds of EASI-FISH (right), separated by the midline (dashed). The three bottom right insets show different 3D views of the crop depicted in I. The left inset shows the crop view from the blue side. The middle inset is a top view. The right inset is a zoom-in of the selected cell and mRNA transcripts detected around it. (J) ΔF/F activity trace of a neuron and its spot counts for 6 genes.

After fixing the brain, we next acquired a volumetric anatomical reference image (Figure 1C). We then gel-embedded the brain using expansion microscopy (ExM) to obtain high imaging quality throughout the entire brain volume and stabilize RNA. We measured mRNA expression levels of 41 genes throughout the brain using multiple rounds of *in situ* hybridization amplified by the hybridization chain reaction (HCR) in the expanded gel (Figure 1D,E; Methods: Expansion-Assisted Iterative Fluorescence *in situ* Hybridization or EASI-FISH, Videos S2 and S3).^12,27^ As expansion microscopy permits sufficient resolution to assign transcripts to densely packed cells, we were able to register these volumes across all 14 rounds of *in situ* hybridization, and back to the live-imaged volumes at single-cell accuracy. To address the challenges accompanying the processing of the samples, including distortions due to fixation, dissection, mounting, and to a lesser extent, expansion, the registration used several intermediate steps. We further applied a number of preprocessing steps to the raw imaging data before all the data was combined for analysis. This data processing pipeline encompasses: (1) Accurate single-cell registration of brain volumes acquired across three modalities (light-sheet functional imaging, two-photon imaging, and imaging of the expanded samples) with new methods to handle the particular types of deformations that occur at each step. (2) High-fidelity segmentation of individual cells across a whole brain volume with a new custom-built distributed version of Cellpose.^22^ (3) Sub-optical resolution limit detection of RNA transcripts, (4) Assignment of RNA transcripts to segmented cells within areas of tightly packed cells using a novel probabilistic algorithm. (5) Inference and demixing of neural calcium signals with a novel cell-segment-initialized Non-negative Matrix Factorization (NMF) algorithm.

This pipeline enabled us to obtain a unique multimodal dataset (Figure 1F,G) that combines functional activity in the form of time-varying calcium signals (Figure 1H and the sagittal plane of brain crop) and RNA transcript counts for 41 genes (Figure 1I and horizontal plane of brain crop) for individual cells (Figure 1J) across the whole brain volume in multiple animals. Our methodology facilitates high-throughput functional and molecular coverage of the whole-brain (Video S4, S5) with over two orders of magnitude more neurons recorded with combined activity and gene expression per animal compared to previous methods.^5,6,8,29^

### Computational methods for accurate and efficient data preprocessing

Our experiments yield high data volume datasets - typically 10+ terabytes per animal - that require multiple preprocessing steps prior to the subsequent analysis procedures. We developed a suite of open-source data processing tools that together solved the challenges presented by this multimodal data (https://github.com/zebrafish-WARP/WARP). The following sections provide a concise overview of these methodologies.

### Whole-brain registrations

Preserving cell identity across multiple modalities in whole-brain volumes presents several challenging registration problems that were solved with single-cell accuracy throughout most of the brain using an open-source package we developed called BigStream^30^ (Figure 2A, Figure S1A-F, and Methods: Image registrations). The recorded volumes can exhibit non-linear deformations, including local stretching, compression, and twisting of the brain created by the multistep sample preparation and imaging pipeline. BigStream implements multiple transformation and optimization schemes to accommodate these deformations across these large brain volumes, including highly flexible piecewise affine transforms able to model a large range of deformation modes, such as low spatial frequency bends and twists, more directly than free-form deformation (Figure 2B, see Methods: Image registrations). Each registration step was validated independently by randomly selecting crops and classifying cells as correctly aligned or misaligned (Methods, Image Registrations). We found cellular correspondence from EASI-FISH to calcium imaging data to be highly accurate (∼80%), sufficient for subsequent analyses (Figure S1A-F).

**Figure 2.**
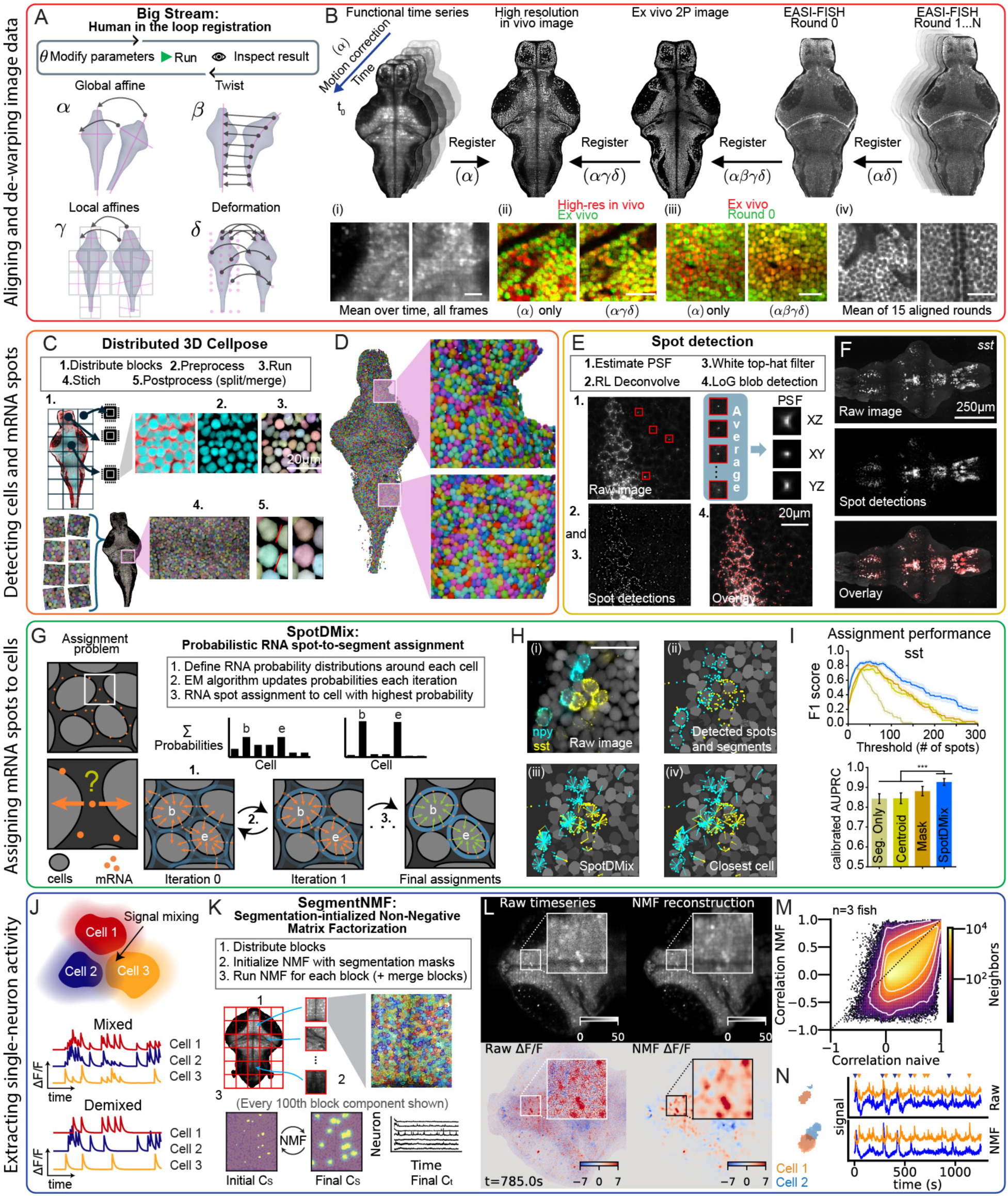
WARPs computational tools for data preprocessing (A) Registration workflow and transform modes available in BigStream with steps ɑ,β,γ,δ applied in (B) (B) Top: Registrations performed at the different steps are indicated with the symbols corresponding to A. Bottom: Validations of registrations. (i) shows the averaged time series of the functional imaging: (ii) the accurate registration between light-sheet high resolution image (red) to *ex vivo* two-photon image (green) with just an affine registration or with all registration steps. (iii) the accurate registration between *ex vivo* two-photon image (red) to the expanded sample (green) with just an affine registration or with all registrations, (iv) the average of registered acquisitions of the EASI-FISH rounds (scale: 20 µm). (C) Nucleus segmentation in large 3D images using distributed Cellpose and post-process corrections. (D) Whole-brain 3D rendering of segmented nuclei. (E) The logic for detecting mRNA transcripts (“spots”). The PSF was calculated to deconvolve the images and detect precise location of transcripts. PSF: Point Spread Function; RL: Richardson–Lucy; LoG: Laplacian of Gaussian filter. (F) Example max projection of the raw image (top), the spots detected (middle), and overlay (bottom) of *sst* mRNA transcripts found in F1 score. (G) The problem of correctly assigning spots to tightly packed cells was solved using SpotDMix^31^, which works by defining probability distributions for RNA transcripts around each cell boundary, then uses the Expectation Maximization (EM) algorithm to update these probabilities and assign accordingly. (H) A spot assignment example using *npy* and *sst* mRNA transcripts: (i) image of the mRNA detected with HCR, (ii) detected spots, (iii) assignments of spots using SpotDMix, (iv) assignments of spots using closest cell boundary (scale: 20 µm). (I) Performance of SpotDMix (dark blue) compared to other naive methods: Segmentation only (spots found within the segmented nucleus (Beige)), Centroid (spots closest to the centroid of the segment (gold)), and Mask (spots closest to the boundary of the segmented mask (orange)). Top: the calibrated F1 score for *sst* based on different thresholds (number of spots). Bottom: Calibrated area under the performance recall curve for *sst*. (Wilcoxon signed-rank test; ns p>0.05; * p<0.05; ** p<0.01; *** p<0.001. Comparisons for *cxn* and *npy* are in Figure S1M, and their statistical comparisons in Figure S1N). (J) Schematic illustrating how the calcium signal of different cells can be mixed as they bleed into other cells. (K) Schematic describing the Non-negative Matrix Factorization (NMF) approach to demix the activity signals. (L) Reconstruction of the signal by SegmentNMF using high-resolution segmentation initialization. Top left: crop and zoom-in of a raw image from the functional time series. Top right: Reconstruction of the signal by SegmentNMF. Bottom left: ΔF/F detected with raw images. Bottom right: ΔF/F detected after SegmentNMF (see Video S7). (M) Correlation between neurons and their close neighbors (r<10 μm) for raw extracted signals, where a cell’s activity is the average of its segment pixels for each time step, and the NMF extracted signals. Signals extracted using NMF have lower correlations within this neighborhood suggesting signal demixing. (N) Example of demixing of the calcium signal of two cells. Top: Raw cell segments and extracted signals. There is clear mixing of signals between the cells. Bottom: NMF allows spatial components initialized with the raw cell segments to grow and partially overlap, enabling demixing of neural signals. Colored arrows indicate activity events for each neuron respectively.

### Nucleus segmentation

We segmented nuclei using Cellpose^28^ to precisely identify the position and a conservative notion of spatial extent of each cell. We used the segmented cells to then assign transcripts and initialize the functional segmentation algorithm for neural activity extraction (see Figure 2C and Methods, Nucleus Instance Segmentation). Specifically, to accommodate the data size and computational load, we designed a distributed version of Cellpose which can handle terabyte scale volumes on a cluster or workstation (Figure 2D) (see the function *cellpose.contrib.distributed_segmentation* in the Cellpose repository: https://github.com/MouseLand/cellpose). We found stitching of both images and segments after distributed computations to be accurate (Figure S1G-I) and validated nucleus detection by measuring the number of nuclear segments found in each brain area, which proved highly consistent across all fish (Figure S1J).

### RNA transcript detection

To extract the precise positions of individual RNA transcripts, we processed EASI-FISH images with a series of filters that algorithmically estimate the optical point spread function, deconvolve the imaged volume, and detect puncta for precise HCR spot localization (Figure 2E,F and Methods: EASI-FISH RNA transcript spot detection). Distributed versions of these functions were implemented to handle the large data volumes (https://github.com/GFleishman/fishspot). We validated our approach by detecting spots for transcripts redundantly probed in multiple rounds and found the number of detected spots to be highly consistent across all brain areas in all fish (Figure S1K).

### RNA transcript to segment assignment

In larval zebrafish, neurons are tightly packed together, with a thin perinuclear cytoplasm. This posed a challenge for determining whether individual RNA transcripts (spots) belong to a segmented cell or one of its neighbors (Figure 2G, left). To address this challenge we developed SpotDMix^31^(https://github.com/Kepser/SpotDMix), a probabilistic method that exploits the fact that a gene typically expresses multiple transcripts in a cell. SpotDMix uses Expectation Maximization (EM)^32^ to assign an RNA transcript to its most likely cell, increasing the probability of assigning a gene transcript to a nucleus that already contains transcripts of the same gene (Figure 2G, right, and Methods: RNA transcript to segment assignment, Video S6). We find that SpotDMix outperforms naive methods that assign transcripts to their closest segment (Figures 2H,I and S1L-N).

### Neural calcium signal extraction and signal demixing

Cells’ calcium fluorescence during activity can extend beyond their physical boundaries due to the microscope’s point-spread function, which results in neighboring cells producing overlapping signals on the imaging sensor (Figure 2J). Non-negative Matrix Factorization (NMF) is an established algorithm for identifying and demixing time-varying neural signals.^33–35^ Taking advantage of the registered high resolution anatomical segmentation from the ExM volume, we built upon the NMF approach to develop SegmentNMF (https://github.com/L-Hess/segmentNMF), an NMF-based calcium fluorescence demixing and extraction approach initialized by the detected anatomical segments of the cells in ExM (Figure 2K, and Methods: Segmentation-initialized NMF). Using the spatial location for each cell provides a strong biological prior that improves cell signal extraction and allows us to track cell identity in all imaging modalities. We found that SegmentNMF denoised the raw imaging input (Figure 2L, Video S7) and reduced correlations between neighboring cells likely caused by signal mixing (Figure 2M,N).

### High-dimensional multimodal datasets

Across all three fish in the main dataset, we segmented a total of 500k+ cells, which comprised neurons, glia, and non-neuronal cells. The multimodal subset of neurons contained ∼238k cells that had both detected gene expression for at least one of the probed genes and were located in one of the imaging planes. The remaining cells in the 500k+ dataset were either *cx43*-positive and therefore likely glia (4.02%), showed no expression of any of the probed genes (39.92%) or were not located in the functionally imaged planes (17.03%). An overview of the neurons of one dataset for a single fish is presented in Figure 3A. The rows (neurons) are ordered by their similarity in neural activity. The experiment was divided in multiple sessions: 1) spontaneous behavior; 2) positional drift; 3) presentation of eight visual stimuli (whole-field visual motion in four directions, dark and bright flash, right and left loom); 4) motor gain adaptation; and 5) futile swimming (Figure S2A; for details, see Methods: Experiment sessions). The experimental setup allowed us to record the activity of most neurons across the brain and the behavioral responses across multiple paradigms (Figure 3A, ‘Activity’). For each of the functionally recorded neurons, the dataset contains RNA transcript counts of the 41 genes probed (Figure 3A, ‘gene expression’), their brain region according to the registered anatomical atlas (Figure 3A, ‘Brain regions’), as well as the behavioral responses of the fish to the presented visual stimuli (Figure 3A, bottom row). An overview of the datasets from the two other recorded fish are shown in Figures S2B and S2C. The probed genes were selected based on the criteria that their composite expression would cover most brain areas, and that their expression partially overlapped, to try to identify cell subtypes with different combinations of genes, together with some genes of particular interest including neuromodulators (Methods: Selection of genes). In addition, we included common genes relevant for neuromodulation (e.g. *net*, *sst*, and *tph2*). We collected another dataset that includes recordings from an additional three fish with a reduced number and different combination of gene probes. These samples were used for validation of our methods but were not further analyzed in this work. They are available as an additional open-access dataset (see Supplemental Information).

**Figure 3.**
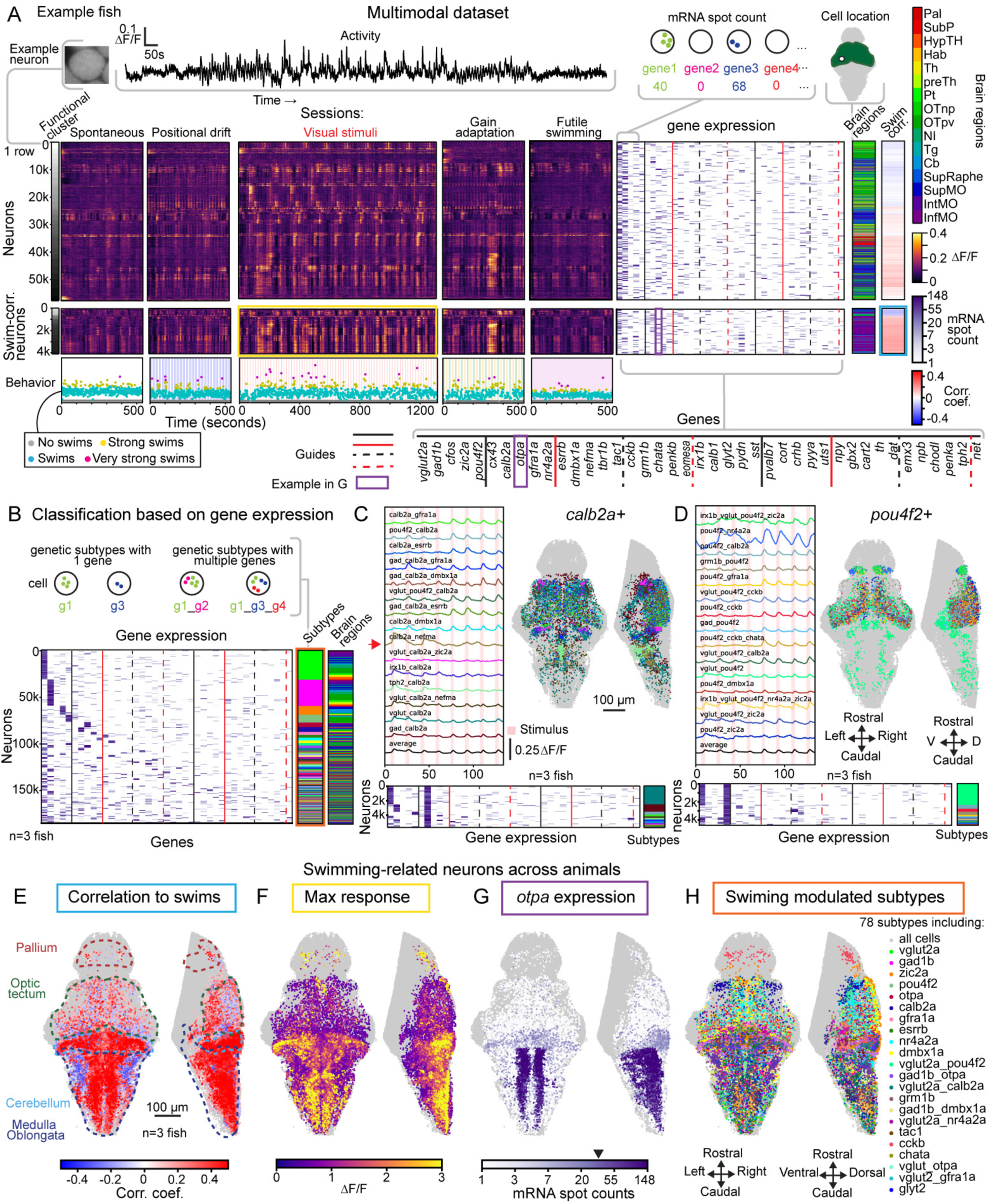
WARP’s high-dimensional multimodal dataset allows for joint analysis of gene expression and activity patterns in the whole brain across multiple animals. (A) Representative single fish dataset. Top: schematic of the different modalities for each single cell. The upper row of matrices and vectors represent the different features, ordered into clusters by similarity of activity during all sessions. The first vector represents the functional cluster (k=30). The next 5 matrices represent neuronal activity traces (ΔF/F) for each of the sessions. The following matrix to the right represents the spot counts of the 41 genes (including *cfos*). The brain region vector represents the location of the neurons based in a brain atlas. The last vector represents the correlation values of the neuronal activity with the swimming behavior (Methods: Correlation to swimming behavior). For clarity, all vectors and matrices were binned every 10 cells (rows) (Methods: Plotting matrices and vectors). The middle row of matrices and vectors show as above, but for cells selected by their correlation to swimming activity (R>0.1 and <-0.1). The bottom row shows the fictive swimming events for each of the sessions. The colored shaded areas in each session represent the visual stimulus presentations. Gray circles = no swims; blue circles = swim; yellow circles = strong swims; magenta circles = very strong swims. The black and red solid and dashed vertical lines are visual guides placed every five genes. Abbreviations: Pal: pallium; SubP: subpallium; HypTH: hypothalamus; Hab: habenula; Th: thalamus; preTh: prethalamus; Pt: pretectum; OTnp: optic tecum neuropil; OTpv: optic tectum periventricular layer; NI: nucleus isthmi; Tg: tegmentum; Cb: cerebellum; SupRaphe: superior raphe; SupMO: superior medulla oblongata; IntMO: intermediate medulla oblongata; InfMO: inferior medulla oblongata. (B) Cells of the 3 samples classified by gene expression. Each molecular subtype had to be found in all 3 fish and adhere to restricted criteria to locate trustworthy replicates (Methods: Anatomical and subtype filtering). The matrix represents spot counts of the 41 genes (including *cfos*). The subtypes vector represents the classification of subtypes based on the gene expression. The brain region vector represents their location based on a brain atlas. All vectors and matrices were binned every 10 cells (rows) for clarity. For categorical variables (genetic cluster, molecular subtypes and brain regions), the representative row was taken from the mode of the 10 corresponding cells. For gene expression, the representative row was taken every 130 cells. The same subtype colormap f is used across panels and figures. (C) Brain map showing cells from the 15 largest molecular subtypes that have positive expression of *calb2a* in common. The top left panel shows the average activity of the neurons of those subtypes. The bottom right vector and matrix represent the subtype classification and spot counts respectively. (D) Brain map showing the cells from the 15 largest subtypes with expression of *pou4f2* and other genes. The top left panel shows the average activity of the neurons of those subtypes. The bottom right vector and matrix represent the subtype classification and spot counts respectively. (E-H). Brain maps showing consensus neurons from the 3 fish that have correlation coefficient values <-0.1 or >0.1 to their respective swimming activity. To be included, neurons of each subtype (shown in H) had to be present in all 3 fish. (E) Correlation coefficient values of the neuron activity traces and the swimming activity. (F) Maximum response activity traces (ΔF/F) of the selected neurons. (G) Brain map showing the spot count values for *otpa* in the selected neurons. Arrowhead indicates threshold (36) to consider a neuron *otpa* positive. (H) Molecular subtypes of the swimming related neurons.

### Classification of cell and molecular subtypes

The quantification of mRNA expression of multiple genes at the resolution of a single cell constitutes features that define its genetic identity. We used these expression profiles to classify neurons into “cell types”, such as *gad1b*+ or *npy*+ cells (when considering only one gene independently of other probed genes), as well as “molecular subtypes”—or, for brevity, “subtypes”—defined by co-expression of multiple genes after binarizing their mRNA transcript counts (Methods: Identifying positive cells). This approach identified molecular subtypes of neurons with varying levels of specificity, depending on how many probed genes each cell expressed. We probed for 41 genes but used 40 genes for molecular subtype analysis as one probed the immediate early gene *cfos*, which we analyze separately (Results: Visual motion and swimming induce *cfos* expression). This led to a range of subtypes, from neurons expressing only a single probed gene (and negative for the other 39 probed genes) to more specific subtypes co-expressing up to 5 probed genes. In total, we identified 332 subtypes, which we denote by their constituent gene names: e.g., a subtype positive only for gene 1 is labeled “gene1”, while a subtype co-expressing gene 1, gene 2, and gene 3 is labeled “gene1_gene2_gene3” (Figure 3B, where we found that there were 40 subtypes defined by 1 gene, 152 defined by 2 genes, 112 by 3, 24 by 4, and 4 subtypes defined by 5 genes). We ensured replication of subtypes across animals by enforcing filtering criteria based on distance between cells and a minimum number of cells, which we compared both within and across fish (Methods: Molecular subtypes).

With the subtypes defined, we next examined the similarity of the activity time series of subtypes sharing a selected gene, focusing on a select few. As an example, we analyzed the 15 subtypes with the most neurons expressing *calb2a*, a gene that encodes *calretinin*, a calcium-binding protein widely used as a neuroanatomical marker (Figure 3C). Some *calb2a* subtypes, particularly those in the hindbrain (e.g. calb2a_nefma, Figure 3C, red arrow), were most responsive to forward visual motion (first stimulus in the visual stimuli session), which elicits swimming. Other *calb2a* subtypes, expressed in the optic tectum (OT) (e.g. calb2a_gfra1a and gad1b_calb2a_dmbx1a), responded most to changes in luminance and to looming stimuli (last four stimuli) (Figure 3C). We also inspected tectal neurons expressing *pou4f2* (Figure 3D), a transcription factor known to be involved in the development and axon guidance of the visual system of fish and mice.^36,37^ Most *pou4f2* subtypes (e.g. pou4f2_gfra1a, pou4f2_cckb_chata) responded to dark and bright flashes, and right and left looms (the last four stimuli in the ‘Visual stimuli’ assay); however, some subtypes (e.g. vglut_pou4f2, pou4f2_zic2a) also responded to both forward and backward visual motion (first and second stimuli) (Figure 3D). These *pou4f2* subtypes may represent distinct subpopulations within the optic tectum whose functions are linked to different visual features. Additional examples of subtypes, including some positive for *vglut2a, gad1b* and *zic2a,* are shown in Figure S3A-F and in Video S8.

In summary, the refinement of cell populations into molecular subtypes provides an insightful classification for comparing and analyzing neuronal activity across fish, experiments and laboratories. Furthermore, our results show that the classification of neurons into functional molecular subtypes facilitates a more brain-wide, targeted interrogation of their role.

### Identification of brain-wide functional neural networks

We next sought to reveal genetically-defined whole-brain functional networks, which we here defined as sets of neurons with enhanced activity correlations with the animal’s behavior. For example, of 332 molecular subtypes, we identified 78 whose calcium activity correlated (R>0.1, higher than 77.52% of neurons) or anticorrelated (R<-0.1, lower than 93.13% of neurons) with swimming, across all three fish (Figure 3A, middle row and Figure S2B-C middle rows; Methods: Correlation to swimming behavior; Finding swimming related neurons; Anatomical and molecular subtype filtering). These swim-related subtypes spanned multiple brain regions, including the pallium, optic tectum, cerebellum and medulla oblongata (Figure 3E-H). The anatomical distribution and correlation coefficients revealed that neurons of the lateral superior medulla oblongata (a subregion of SupMO termed L-MO) were anticorrelated with swim strength, consistent with its known function^25,38^ while many neurons in the cerebellum and the medial hindbrain were strongly correlated with swim strength (Figure 3E; maximum intensity (of absolute values) shown; minimum intensity in Figure S2D highlighting negatively-motor-correlated neurons). While collectively correlated with swimming, these cells did exhibit variability in their activity profiles (Figure 3A, yellow box) and response intensities (Figure 3F). We further note that many swim-related neurons, particularly the ones located in the medial hindbrain, showed high expression (number of mRNA spots) of the transcription factor *otpa* (Figure 3G) suggesting a possible role for this gene in shaping swimming behavior.

In summary, this example illustrates how WARP, through analyzing brain-wide behavior-correlated neuronal assemblies in terms of precisely defined molecular subtypes, enabled us to map a diverse distribution of neuronal subpopulations that were consistently observed across fish (Figure 3H, Video S9). WARP thus facilitates a high-throughput investigation into the relationship between the molecular and anatomical properties of brain-wide neuronal populations and how their activity patterns relate to specific behaviors and each other.

### Brain-wide gene expression patterns reveal stereotypical cell subpopulations

Having established the framework of WARP, we next investigated the relationships between brain-wide gene expression, co-expression patterns, and stimulus response patterns. We first assessed how well our gene list covered 16 major anatomically defined brain regions^39^ (Figure 4A). For the following analyses, we include neurons outside of light-sheet imaging planes for anatomical analyses with a total of ∼306k cells, extending the set of 238k cells also including functional data. As above, included cells were selected based on sufficient RNA expression to be considered positive for at least one of the 41 (including *cfos*) genes probed (detected RNA transcripts above a threshold; see Methods: Identifying positive cells and Data S1. Neural activity analyses below are based on the subset of 238k cells). Their total spatial distribution spanned regions across the entire brain. The hindbrain subdivisions, superior (SupMO), intermediate (IntMO) and inferior (InfMO) medulla oblongata, showed the densest coverage of probed genes. By contrast, the habenula, tectal neuropil, and superior raphe had the lowest gene coverage (Figure 4A). Importantly, gene expression patterns were consistent across animals (Figure 4A, Figure S4, Figure S5 and Video S10) and in line with previous studies where available.^18,40–46^

**Figure 4.**
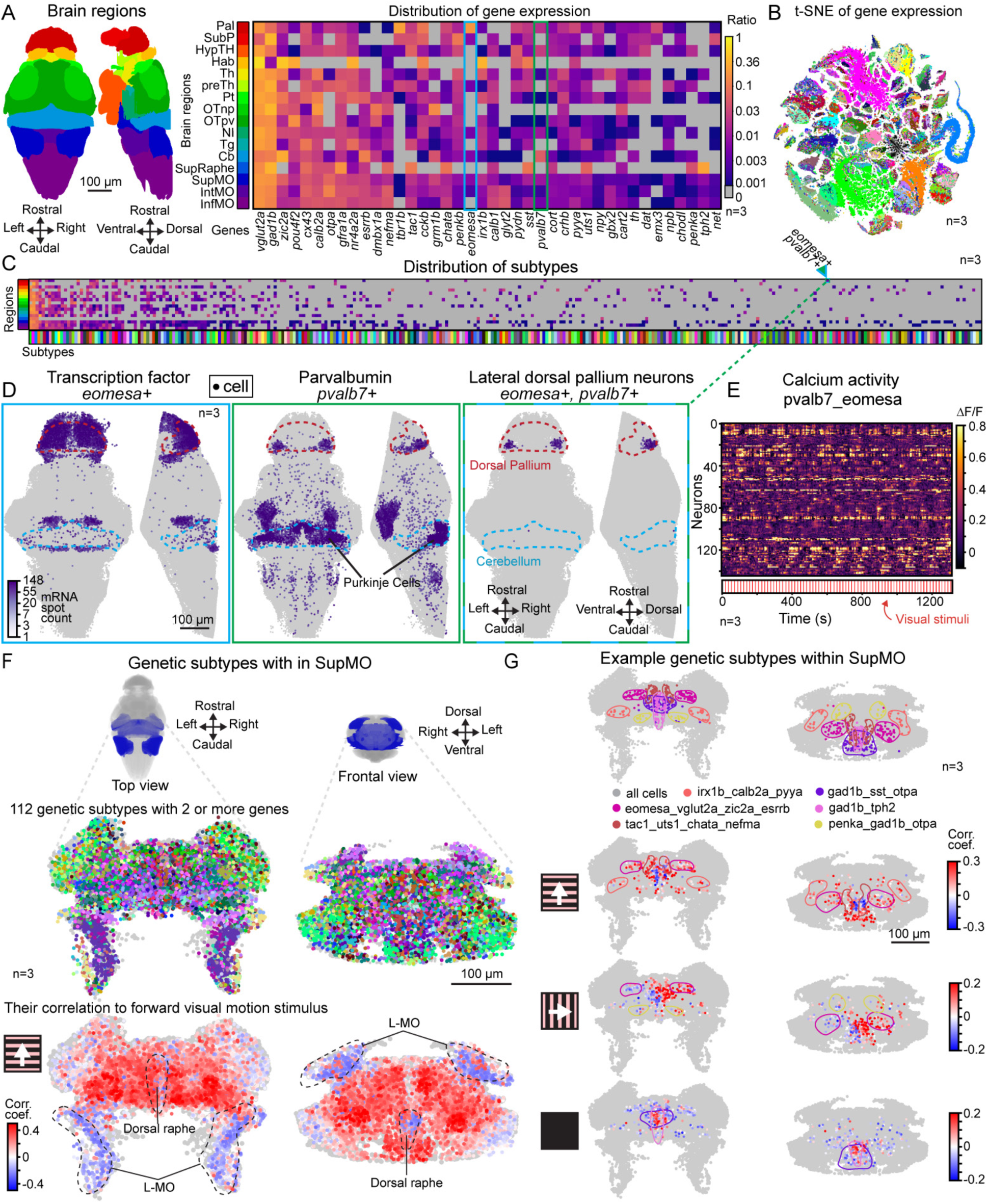
Observed brain-wide spatio-transcriptomic expression patterns reveal stereotypical cell subpopulations that are consistent across samples. (A) Left: Common brain regions of interest defined by a reference atlas (mapZebrain^39^). Right: Average percentage of positive cells for the 40 (excluding *cfos*) genes within each of the brain regions (n=3). (B) t-SNE visualization of a 2-dimensional embedding of the gene expression in 306,108 cells from all 3 fish. Colors represent the 332 subtypes. This visualization is also shown with labels per fish (Figure S6B), spot counts (Figure S6C), and brain region (Figure S6D). (C) Average percentage of the 332 subtypes across 16 brain regions. A complete version of this matrix is in Figure S7A. (D) Distribution of cells expressing *eomesa* (left), *pvalb7* (middle), and both *eomesa* and *pvalb7* (right). Cells from all 3 fish are included. (E) The calcium activity of the cells expressing both *eomesa* and *pvalb7* in D. (F) Distribution of molecular subtypes in the SupMO and their correlation to forward visual motion. Top: Reference top and frontal view of the SupMO. Middle: Distribution of the 112 molecular subtypes in the SupMO that are positive for 2 or more genes. Bottom: Correlation coefficients to the forward visual motion stimulus presentation of the above cells. Higher absolute values are plotted on top. Cells from all 3 fish are included. (G) Examples of molecular subtypes selected by their higher locality index and a higher-than-average inner correlation. Top: Distribution and contour of subtype location. Bottom: Correlation coefficients to the forward visual motion, the rightward visual motion and the dark flash. Cells from all 3 fish are included. Abbreviations: Pal: Dorsal Pallium; SubP: Subpallium; HypTh: Hypothalamus; Hab: Habenula; Th: Dorsal Thalamus; preTh: Prethalamus; Pt: Pretectum; OTnp: Optic tectum neuropil; OTpv: Optic tectum periventricular layer; NI: Nucleus Isthmi; Tg: Tegmentum; Cb: Cerebellum. SupRaphe: Superior dorsal raphe; SupMO: Superior Medulla Oblongata; IntOM: Intermediate Medulla Oblongata; InfMO: Inferior Medulla Oblongata.

Leveraging the enhanced multiplexed nature of WARP, we next analyzed transcript co-expression in single neurons (Figure 4B), and their distribution across the brain (Figure 4C). We observed expected relationships, such as *dat*+ or *net*+ neurons co-expressing *th1*,^47,48^ as well as lesser-known co-expression patterns such as *th1* with *calb2a* or *cort*^49^ (see Figure S6A for these and other examples). Using our single-cell, whole-brain dataset, we then identified subtype spatial distributions, such as the *eomesa*+/*tbr1b+* or the *eomesa*+/*pvalb7*+ expressing neurons in the telencephalon.^50^ The expression pattern of *eomesa*, a transcription factor associated with telencephalic development in both mammals and teleosts,^51–53^ was observed in telencephalic neurons (Figure 4D, left). Among the other genes identified in the telencephalon, *pvalb7*+ (parvalbumin) neurons were located in the lateral edges of the dorsal pallium (Figure 4D, middle). Most of these *pvalb7*+ neurons also expressed *eomesa* (Figure 4D, right; this co-expression has also been confirmed in scRNA-seq data^50^), indicating that they belong to a neural population in the dorsolateral pallium that previous studies suggest is homologous to the mammalian hippocampus.^50,54,55^ Notably, although this structure is in its early development at 6 days post fertilization (dpf),^56–58^ these *pvalb7*+ neurons were already active, showing task-related calcium dynamics (Figure 4E), indicating functional activity within the first week of life. Additionally, *pvalb7*+ cells were also found in the cerebellum, where they correspond to the well-known Purkinje cells.^43^ Our spatial transcript analysis thus revealed both expected observations and new insights into the distribution of subtype populations across the brain. While we cannot exhaustively describe all subtypes here and have to focus on a few examples, this open-access dataset provides a valuable resource to investigate specific function-gene-anatomy-behavior relationships.

### Molecular subtypes exhibit diverse but specific distributions

Compared to single genes, the molecular subtypes defined above provide a greater level of detail, revealing a variety of population sizes and distributions. Some subtypes were widely expressed across the brain, in particular ones that were only positive for one of the common neurotransmitters or transcription factors (e.g. *vglut2a*, *gad1b*, *zic2a*), while those that were intersections of genes were represented by smaller, more localized populations (e.g. penka_gad_opta and gad_uts1_nr4a2a_gbx2, Figures S7 and S8). The hindbrain contained the greatest number of subtypes (129 in SupMO, 99, IntMO and 80 in InfMO), followed by the optic tectum (79). Within brain regions, different neuronal subtypes often showed distinct spatial distributions that were not apparent without the combined genetic information. For example, in the SupMO some subtypes were tightly packed (e.g. tac1_uts1_chata_nefma; Figure S8) while others were distributed across large regions (e.g. gad_otpa; Figure S7A and Figure S8). Furthermore, compact clusters took on different shapes, which could be approximately spherical (e.g. eomesa_vglut2a_zic2a_esrrb; Figure S8) or columnar (e.g. vglut2a_otpa; Figure S8). These findings underscore how, through detailed characterization of the genetic profiles and spatial context of the cells, it is possible to further define molecular features of classical anatomical regions, refining existing subregions^12,59–62^ and defining new ones, which can then be used to more accurately localize substructures within specific brain areas.^63^

We asked if the tendency for cells of a given subtype to be clustered or spread out varied by brain region and if such differences could reveal further details about the spatial organization of the subtype. We first calculated a locality index for each subtype within each anatomical region (Methods: Spatial locality index and inner correlation for subtypes in brain regions), revealing that the pallium, thalamus and SupMO were regions with the most spatially compact subtypes, while the optic tectum had the least, likely due to its layered organization (Figure S9A). We next asked if the correlations of the activity of subtypes depended on the degree to which they were tightly or loosely packed in space. Interestingly, the average pairwise activity correlation of neurons within a subtype (“inner correlations”, Figure S9B) increased with locality index (Figure S9D-F; R=0.339), suggesting that spatially tighter aggregates of the same molecular subtype are more likely to constitute a more activity-synchronized functional neuronal group.

To explore more deeply how the spatial distribution of subtypes might participate in functional networks (i.e. groups of cells with enhanced activity correlations), we examined the subtypes found in the SupMO region. This region contains the largest number of subtypes and has an established role in swimming and the optomotor response^17,20,64^ (Figure 4F, top and middle). We first analyzed the correlation between the activity of the 112 subtypes and the forward visual motion stimulus. A broad area of the SupMO showed high correlations between its neural activity and this stimulus time series (Figure 4F, bottom), while specific regions like the lateral medulla oblongata (L-MO) and the dorsal raphe showed negative correlations. The SupMO had the second highest number of subtypes with both locality index and inner correlations above the mean for the entire brain (Figure S9C and S9F). To examine if these molecular subtypes coordinate their activity in a specific way, we selected six of these molecular subtypes with higher-than-average spatial locality index and inner correlation and plotted the correlation coefficients of their neurons’ activity in response to forward and sideways visual motion, and dark flashes. Five subtypes showed higher correlations with the forward visual motion, while another, gad_tph2, was negatively correlated, suggesting that this subtype receives inhibition during optomotor responses.^65,66^ Interestingly, this pattern was reversed with dark flashes, suggesting that these cells were also part of the dimming sensitive *tph2* neurons.^67^ Furthermore, during sideways visual motion stimuli, we observed a clear lateral asymmetry in some of the subtypes, with one side showing high correlations and the other negative correlations (Figure 4G). More subtypes are shown in Figure S9G.

Taken together, this analysis demonstrates how neighboring neuronal populations with different genetic profiles can be recruited for similar or distinct functional roles, and, by integrating functional, genetic, and anatomical information, provides the substrate for a highly specific and comprehensive functional-genetic-anatomical brain atlas.

### High-throughput discovery of visually responsive subtypes throughout the brain

Using the framework and approaches developed in the previous sections, we expanded our analysis across the entire brain to determine which molecular subtypes were active when the different visual stimuli were presented (Figure 5A).^17,20,24,68–73^ Of note, the processing of visual stimuli trigger both sensory and motor responses, as well as engage neuromodulator populations.^64,66,74^

**Figure 5.**
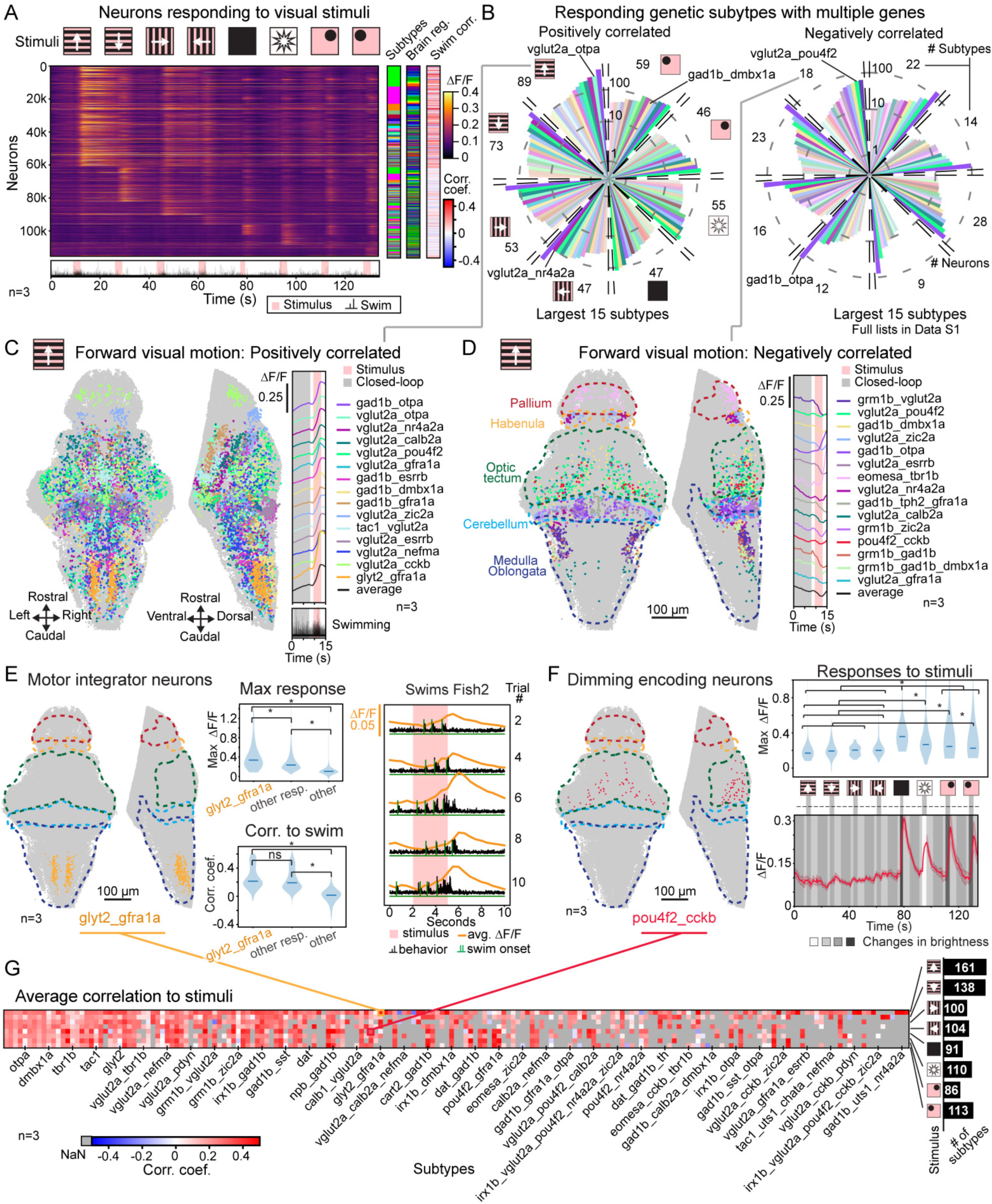
Over a hundred molecular subtypes are involved in visual stimulus responses. (A) Average trial activity trace of the neurons that responded to the eight visual stimuli. Their molecular subtype, brain region, and correlation to swimming is shown on the right. Neurons from all 3 fish are included. At the bottom the behavioral (swim vigor) responses of all 3 fish are overlaid. (B) The 15 largest (highest number of neurons, averaged across the 3 fish) molecular subtypes with 2 or more genes involved in the responses to the visual stimuli. The number in each section is the total number of subtypes (with multiple genes) that were consistent across experiments (the rest of the responsive subtypes can be found in Data S1). The left side shows results for positively correlated responses to the stimuli, the right side for negatively correlated responses. (C) The 15 largest multi-gene subtypes positively correlated with forward visual motion. Left: Top view of these subtypes in their anatomical location. Middle: lateral view. Right: Average activity trace of each subtype. The average is calculated from the neurons of the 3 fish. (D) The 15 largest multi-gene subtypes negatively correlated with forward visual motion. Left: Top view of these subtypes in their anatomical location. Middle: lateral view. Right: Average activity trace of each subtype. The average is calculated from neurons of all 3 fish. (E) Motor integrator neurons that co-express *glyt2a* and *gfra1a*. Left: anatomical location of glyt2a_gfra1a cells positively correlated to forward visual motion. Neurons from the 3 fish are included. Middle-top: Maximum response during the stimulus presentation of the glyt2_gfra1a cells, the rest of forward visual motion responding cells and the non-responding cells in the InfMO (Kruskal-Wallis test, p<0.001, followed by Dunn’s test with Bonferroni correction. All paired comparisons indicated with the dendrogram with a * have p<0.05). Middle-bottom: correlation to swimming of the glyt2_gfra1a cells, the rest of forward visual motion responding cells, and the non-responding cells in the InfMO (Kruskal-Wallis test, p<0.001, followed by Dunn’s test with Bonferroni correction. All paired comparisons indicated with the dendrogram with a * have p-values under 0.05). Right: the average calcium trace of glyt2a_gfra1a cells for Fish2 at 5 of the presentations of the forward visual motion stimulus. (F) pou4f2_cckb dimming light responsive neurons. Left: pou4f2_cckb cells positively correlated to a dark flash are located in the optic tectum. Neurons from all 3 fish are included. Right-top: Maximum response of these pou4f2_cckb cells during the stimulus presentations (Kruskal-Wallis test, p<0.001, followed by Dunn’s test with Bonferroni correction. All paired comparisons indicated with the dendrogram with a * have p-values under 0.05). Right-bottom: Average activity trace of the pou4f2_cckb cells and the changes in luminance caused by the stimulus presentations. The average is calculated from neurons of all 3 fish. Shaded areas show ± SEM. (G) Matrix with the correlation coefficient average for each of the stimuli. Each row represents one of the stimuli presented, each column represents one of the defined subtypes. Subtypes are ordered as in Figure 4D.

Within neurons that correlated to each of the presented stimuli, we found 183 molecular subtypes that consistently responded across all three animals. Among the largest subtypes expressing two or more genes and showing positive or negative correlations to each of the stimuli (Figure 5B, Video S11; Data S1) there were five frequent subtypes: gad1b_otpa, vglut2a_pou4f2, vglut2a_nr4a2a, vglut2a_calb2a, and gad1b_dmbx1a. We next selected 15 of the largest subtypes that responded to the forward visual motion stimulus and examined their anatomical locations and stimulus-response profiles (Figure 5C). These included the vglut2a_cckb (cholecystokininergic) subtype in the dorsal pallium and hypothalamus; vglut2a_pou4f2 and gad1b_dmbx1a subtypes in the optic tectum; gad1b_otpa and vglut2a_nefma subtypes throughout the hindbrain and the glyt2_gfra1a (glycinergic) subtype in the caudal hindbrain. Subtypes negatively correlated to the stimulus (Figure 5D) included the gad1b_tph2_gfra1a in the dorsal raphe, and grm1b_vlugt2a, grm1b_gad1b and grm1b_gad1b_dmbx1 subtypes in the lateral hindbrain (L-MO). These particular subtypes likely correspond to reported neurons in this area that are known to inhibit swimming.^25,38^ Subtype analysis of other visual stimuli show that there was a wide variety, distributed across multiple brain regions (Figure S10).

Closer inspection of several stimulus-responding subtypes revealed specific functional roles. For example, a group of glyt2_gfra1a neurons in the caudal hindbrain responded particularly strongly during forward visual motion and ramped up continuously during swimming (Figure 5E). As glycine-expressing inhibitory neurons, they appear to integrate motor responses elicited by the stimulus, suggesting a role in providing negative feedback after successful execution of swimming.^75,76^ Another example of specific functional roles are the dimming sensitive pou4f2_cckb neurons in the optic tectum that responded to dark flash and looming stimuli (Figure 5F). This analysis illustrates how combining both neuronal activity and gene expression can refine our understanding of the specific location and functional roles of molecular subtypes (Figure 5G).

The examples described above illustrate only a small fraction of the total number of such functional-genetic-anatomical subpopulations. To generate a comprehensive set of subpopulations classified by these parameters, we categorized neurons according to their correlations to the 8 visual stimuli, their gene expression profile as one of the 329 molecular subtypes (excluding those with *cx43*), and location in one of the 16 main brain areas. After applying a strict set of inclusion criteria (Methods: Subtypes correlation to stimuli matrix), but allowing for partial overlap in cases where neurons respond to multiple stimuli, and enforcing consistency across the three fish, this method yielded a total of 2,256 subpopulations involved in the processing of these ethologically relevant cues and resulting behavioral outputs, producing an unprecedented functional-genetic-anatomy based description of the brain (average number of neurons = 1,009, SD = 2,137, min = 109, max = 22,278. See Data S1).

### Visual motion and swimming induce *cfos* expression

To investigate whether the stimuli and evoked swimming behavior induced molecular changes in the responding neurons, we probed for the expression of the immediate early gene *cfos*, a widely used molecular marker for neuronal activity^77,78^ (Methods: *cfos* labeling and analysis; Figure S11A). We found that *cfos+* neurons exhibited stronger responses compared to *cfos-* neurons, had larger activity fluctuations, and showed higher correlations to the visual stimuli and to swimming (Figure S11B). Neurons expressing *cfos* also correlated more to the forward visual motion stimulus, and to a lesser extent, to visual motion in other directions and the looming stimuli, consistent with forward motion eliciting the strongest overall brain activity due to its potent swim-driving properties (Figure S11C). Spatially, most *cfos*+ neurons were located in the hypothalamus, rostral hindbrain tectum, and pretectum (Figure S11A and D). These neurons mostly co-expressed *vglut2a* and *gad1b*, but other subtypes were also identified expressing the stress responsive neuropeptides urocortin (*uts1*) and corticotropin-releasing hormone (*crhb*) (Figure S11E). These findings demonstrate that WARP can be used to probe for genes informative of physiological changes and molecular cascades in combination with time-resolved brain-wide neuronal activity measurements.

### Genetically defined cell types exhibit heightened activity correlation across the brain

We next sought a more rigorous quantification of how neuronal gene expression relates to activity. This question speaks to a central hypothesis in systems neuroscience: that a neuron’s molecular identity constrains its functional role. Previous studies have either inferred such gene–function links indirectly^79,80^ or probed them within individual neural circuits.^81,8,5^ However, these anatomically restricted approaches have not established the extent to which molecular identity predicts function across the diverse cellular environments of the whole brain. As our dataset jointly quantifies both modalities brain-wide across dozens of genes, it enabled a systematic, brain-wide test of this hypothesis.

Specifically, we asked whether neurons sharing expression of any one of our probed genes exhibit more similar activity than would be expected given their local surroundings. To quantify this, we devised the Local Correlation Difference (LCD), which compares the activity correlation between two neurons expressing the same gene to a local baseline (Figure 6A). This baseline is defined by the correlation between one of these neurons (the reference neuron) and nearby neurons that do not express the gene within a 15 µm radius. By subtracting this neighborhood baseline correlation, the LCD isolates the association between gene identity and functional coupling from broader spatial or regional correlation structure (Methods: Calculation of Local Correlation Difference).

**Figure 6.**
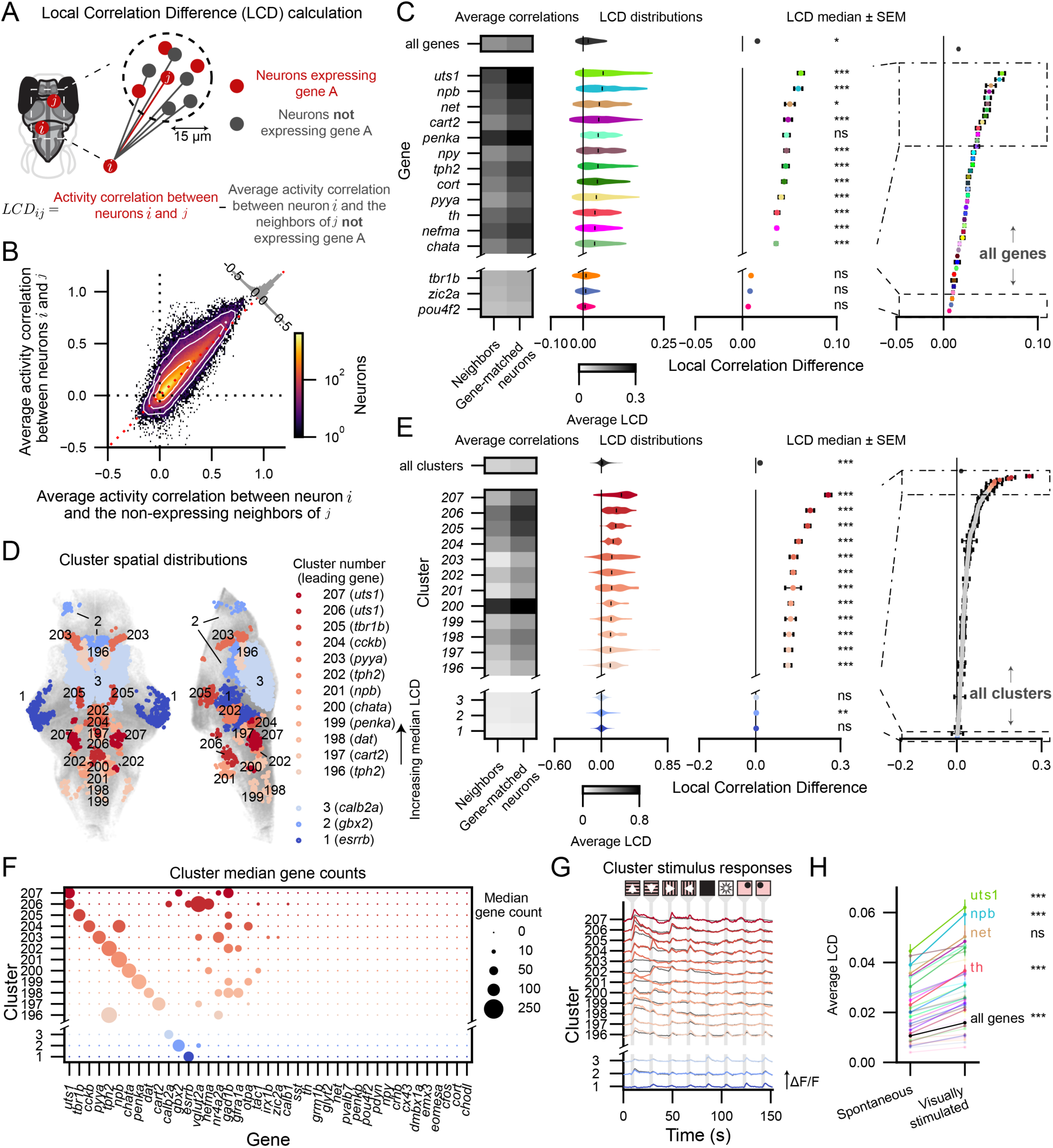
Neural activity is more correlated between neurons of the same transcriptomic type and identified correlation distributions are more extreme in spatially constrained clusters. (A) The Local Correlation Difference (LCD) quantifies how strongly neurons that express the same gene correlate in their activity relative to the local neighborhood. For each gene-matched neuron pair, the LCD is defined as the difference between the activity correlation of the two neurons (red) and the average correlation between the inspected neuron *i* and the non-expressing neighbors of the reference neuron *j* (grey) within a radius of 15 μm with respect to the location of the reference neuron. (B) Joint density of within–gene-type correlations (ordinate) and remaining-neighbor correlations (abscissa) for all genes. Higher within-gene-type correlations, indicated by a larger density-mass above the identity line, produce a positive-shifted LCD distribution. White lines indicate equal-density contours. Inset histogram shows the difference between the ordinate and abscissa and thus indicate LCD values. (C) Local gene-type-specific coordination varies by gene but is significant for most. First column: mean correlation with same-gene-type neighbors vs. remaining neighbors for the 12 genes with highest median LCD and the 3 with the lowest; second column: LCD distributions for the same genes; third column: median ± SEM and significance of LCDs for these genes; fourth column: same statistics for all measured genes. All columns are ordered according to the median LCD. Full LCD distributions in Figure S12B. (two-sided permutation test, BH-corrected across genes; ns p>0.05; * p<0.05; ** p<0.01; *** p<0.001; across all genes, p=0.184 (Figure S12B) p<ε; per-gene values in Data S1). (D) For each gene, spatially constrained clustering groups neurons based on their anatomical locations (see Methods: Spatial clustering of Correlation Differences). This identifies spatially localized clusters of neurons with particularly high or low LCDs that are confined to specific brain regions and often co-express multiple genes. Examples of the 12 clusters with the highest and the 3 clusters with the lowest median LCDs are shown together with their spatial locations. (E) LCD distributions for each cluster, columns as in panel C. Colors match panel D. (Statistics as in panel C; two-sided permutation test, BH-corrected across clusters; across all clusters p<ε; per-cluster values in Data S1). (F) Gene expression profiles per cluster. Dot size indicates the median transcript count for each gene within a cluster. Colors as in panel D. The x-axis is ordered according to the leading genes of each cluster. (G) Average neural responses to the full stimulus train for all neurons in each cluster. Colors as in panel D. Grey lines indicate the average neural responses to all neighboring neurons of that cluster not expressing the same cluster-specific gene. Shaded areas indicate ± SEM. (H) Gene-type-specific LCDs are enhanced by visual stimulation in most genes. 5 genes with the largest increase between the spontaneous and visually stimulated paradigm are highlighted. (Two-sided Wilcoxon signed-rank test, Bonferroni-corrected; across all genes p<ε; per-gene values in Data S1).

To reduce noise at the neuron-pair level, we also computed a smoothed LCD estimate by averaging LCD values across all gene-matched pairs whose reference neuron lay within a 20 µm neighborhood (Methods: Calculation of Local Correlation Difference; Figure S12A). This local LCD average per neuron provided a more stable measure of gene-specific coupling, and used it for subsequent calculations. We performed LCD analysis using broad cell-types (expressing versus not expressing single genes; excluding *cfos*, and *cx43*) rather than the finer multi-gene molecular subtypes defined previously as many subtypes contained too few nearby neurons to yield reliable neighborhood averages.

Consistent with earlier gene-function studies,^7,82,5^ neurons generally showed higher average activity correlations with gene-matched neighbors than with their non-matched neighbors (Figure 6B). LCD distributions differed systematically across genes (Figure 6C, Figure S12B), with some—such as *npb*, *net*, and *uts1*—showing particularly strong within-group activity correlation. For the majority of genes (30/39), the median LCD was significantly greater than zero relative to a null distribution obtained by permutating gene labels (Methods: Permutation-based null model for Local Correlation Differences). This finding indicates that neurons that share expression of at least one gene tended to have more synchronized activity with each other than with other nearby neurons. The pervasiveness and robustness of this effect across genes supports the idea that a cell’s gene expression profile is tightly linked to its functional role across molecular cell types.

While gene expression did correlate to neural activity in local neighborhoods, the LCD values for most genes were not uniformly distributed across the brain, resulting in broad LCD distributions (Figure 6C, violin plots). Within these distributions, we found spatially coherent groups of neurons with distinct levels of gene-specific activity correlation (Figure S12C shows the LCD distributions for *tbr1b* and *uts1*). To identify these distinct subgroups, we clustered each gene’s neurons based on their location in the brain using Leiden-based graph clustering^83^ (Methods: Spatial clustering of Local Correlation Differences) and identified 207 spatially localized clusters across all measured genes. As expected, these clusters exhibited narrower LCD distributions and more extreme median values than the full gene populations (Figure 6D,E; the 12 clusters with the highest and 3 clusters with the lowest median LCDs are shown).

Many clusters co-expressed multiple genes (Figure 6F) and showed distinct responses to the visual stimuli (Figure 6G). For example, most neurons from the cluster with the largest median LCD (spatially localized cluster 207) co-expressed *uts1*, *nr4a2a*, *gad1b*, and *gbx2*, indicating that it represents a distinct genetic subtype. Functionally, this cluster responded strongly to forward and turning visual motion stimuli, but responses were attenuated during backward visual motion and looming stimuli. Together with its location in the intermediate MO, this suggests a role in processing swimming-related behavior. Because LCD aggregates correlations over the full time window, we further decomposed the LCD into per-time-step contributions (Figures S12D and S12E). Plotting these for cluster 207 shows that the forward, left, and right visual motion stimulus periods contribute most to the total LCD. Consistently, correlating these time-resolved LCD contributions with swimming behavior revealed elevated contributions during swim attempts (Figure S12F). Thus, cluster 207 exhibits behavior-linked coordination that is stronger than that of nearby neurons not expressing *uts1*.

More generally, decomposing the LCD over time (Figures S12D and S12E) provides insight into gene-type-specific coordination to specific stimulus epochs (Figure S12F). Notably, while some clusters show coupling to behavior (e.g. spatially localized clusters 207, 206, and 203), most do not, suggesting additional roles in sensory processing, internal state modulation, or other computations beyond processing of motor-related behavior.

We further observe that cluster 205 expresses *tbr1b*, which has one of the lowest median LCDs across its full population (Figure 6C). Consistent with earlier results (e.g., Figs. 4 and 5) and studies^84^, this further illustrates how incorporating spatial information can reveal distinct functional subpopulations within a given genetic class.

Applying the LCD calculations to periods of spontaneous activity (i.e., without external stimulation) yielded significant positive median LCD values for the majority of genes (28/39) (Figure S12B), and under visual stimulation, the LCD significantly increased for almost all (35/39, Figure 6H). These findings suggest that gene-defined neuronal groups maintain functional coupling even in the absence of sensory input, appearing to form intrinsically coherent ensembles whose coupling is further structured during sensory processing.

In summary, our data uncover a link between gene expression and function, whereby common gene expression constrains coordinated activity, mostly elevating coactivation. This principle generalizes to most genes measured across the entire brain, is already present in spontaneous activity, and enhanced during visual stimulation.

## Discussion

This study unifies two major approaches to understanding the brain that are typically studied separately: the cell biology of the nervous system—which describes molecular and genetic cell identity—and whole-brain dynamics, which captures the coordinated activity of large neural networks during behavior. By connecting the molecular, anatomical, and functional domains across the whole-brain at the single cell scale within individuals, our work enables new forms of insight into how the molecular properties of cells shape systems-level computation.

By integrating whole-brain activity imaging, optimized multiplexed FISH,^12,27^ and a suite of data processing and analysis algorithms, we identified the neural dynamics of over 2,000 genetically, anatomically, and functionally distinct subpopulations in single animals; found that activity of neurons of the same cell type tend to be more correlated; that specific subtypes responding to visual stimuli are replicated in different animals; that results were consistent across animals and, where available, with previous findings.^50,76,84,85^ The resulting datasets provide a resource for future discovery by the scientific community, facilitating more detailed computational neuroscience analyses,^86–89^ and enable further community analysis of the recorded paradigms not analyzed in this work. Additionally, these datasets can serve as a tool for educational purposes in the context of neuroscientific discovery and AI methods for biology.^90–93^

WARP provides several significant advancements for the neuroscientific community. For example, it facilitates the high-throughput investigation of the molecular basis of neuronal activity by capturing the dynamics of cells alongside their molecular characteristics. This allows one to correlate observed activity patterns with gene expression across the whole brain as shown in multiple ways in this study. This property furthermore can improve cross-animal alignment of experiments to probabilistically identify small groups of neurons across different animals based on their location in the brain, their responses to stimuli and behavior, and the genes they express.^94^ Furthermore, by mapping activity to gene expression at scale, WARP enables scientists to determine useful marker genes at high throughput for groups of functionally identified neurons almost anywhere in the brain. This property can be leveraged to develop targeted opto-or chemogenetic experiments to determine their causal role for brain dynamics and behavior. Finally, novel applications of the technology may extend to questions on neural plasticity, development, and the effects of environmental perturbations.

Beyond these direct applications, WARP has broader implications for multimodal brain mapping and modeling. By enabling neuron-type identification, it allows atlases^18,39,95^ of anatomy, gene expression, and function to incorporate single-cell, multimodal data. This further extends towards computational and AI models that will be able to use the type-labeled neuronal activity traces to probabilistically assign neuronal types across animals. In this way, WARP’s large-scale multimodal output can be leveraged to assemble arbitrarily large datasets to build increasingly detailed models of brain dynamics.^96,97^ WARP can also advance biologically grounded modeling by addressing a key limitation of current approaches: most models are built from activity and anatomy only,^98–100^ while the minority that incorporate molecular information typically rely on dense labeling restricted to single regions,^84^ or small numbers of molecularly-defined cell types that do not cover the entire brain. With brain-wide, comprehensive, neuron-type identification, WARP can reduce the amount of unaccounted heterogeneity and can thereby increase the explanatory power of fitted models. On top of this, with parallel efforts into recording both function and the full connectome in zebrafish larvae,^100,101^ a comprehensive dataset of functional, structural, and genetic properties of a vertebrate brain is soon within reach. We therefore anticipate that WARP will accelerate mechanistic discovery of brain computation and play a key role in the AI revolution in systems neuroscience.

The ease and accuracy with which WARP allows the comparison of specific subpopulations across animals can also be extended to other species, such as *Danionella*,^102,103^ Mexican cavefish,^104^ and tadpoles,^105^ for which these brain-wide methods can all be applied. While current imaging methods for mice and larger vertebrates are limited to subregions of the brain, our protocol can also be used in thick tissue samples of these animals. Thereby, WARP would extend this comprehensive approach to these models and facilitate interspecies comparisons, elucidating the evolutionary conservation or divergence of specific neuronal and glia cell types.

WARP’s flexible nature provides a platform for further development of its throughput, to record additional modalities, and to acquire additional cell information. While our RNA visualization method is currently limited to dozens of genes, optimizations to use combinatorial techniques^11,106^ could allow probing hundreds or thousands of genes. WARP can be integrated with molecular sensors for voltage,^107^ neurotransmitters,^108,109^ neuromodulators,^110,111^ intracellular messengers,^112^ and other signals, paving the way for an ever-more comprehensive mechanistic understanding of cellular and network dynamics. Additionally, methods for light-based connecomics^113^ can be further developed to be compatible with WARP, enabling the integration of neuronal morphology and electrical and chemical connectivity.

Finally, WARP is not limited to the brain; it can be applied to any tissue or organ—or even the entire body—enabling joint functional and molecular analyses across the whole organism. Its versatility allows for imaging calcium activity in non-neural tissues such as the liver or heart,^114^ followed by identification of the corresponding cell-molecular profiles. This compatibility with other live imaging modalities, including metabolic indicators, offers a powerful platform for whole-body cellular dynamics.

### Limitations and future directions

The integration of neuronal function and RNA profiles across an entire brain presents opportunities as well as challenges and limitations, both on the experimental and the computational side. First, imaging large volumes, or areas that may be harder to image at high resolution, can pose a challenge for cell-accurate registration. Other imaging modalities like multi-photon microscopy,^115^ the use of adaptive optics (hardware^116,117^ or computational^118,119^), or improved neural activity indicators could improve the imaging quality and thereby these registrations. Second, the performance of our probabilistic transcript assignment algorithm, SpotDMix,^31^ improves with a higher number of detected transcripts around gene-expressing cells. However, some genes, such as receptors, tend to exhibit low RNA expression, which increases uncertainty. Increased expansion of the tissue, in combination with light-based connectomics,^113^ would further improve transcript localization and their assignment. Additionally, the molecular subtypes defined in this study are limited to our selected list of genes, highlighting the need for thoughtful gene selection tailored to specific research questions, as well as future adaptation to higher-throughput techniques that can still be used in thick tissue samples like cycleHCR^11^ and weMERFISH.^120^ Finally, the computational methods and analysis presented here serve as a foundation for further exploration and could be expanded using more advanced machine learning methods,^121^ or multi-view approaches^122^ that integrate multiple cell modalities^123–125^ for additional biological insights.

## Resource Availability

### Lead Contact

Further information and requests for resources and reagents should be directed to Misha B. Ahrens

### Materials Availability

This study did not generate new unique reagents.

### Data and Code Availability

- All data generated in this study have been deposited at figshare (https://figshare.com/s/d1d19b105c4f74865c32). Datasets include:

- Activity traces and xyz locations for segmented cells for each fish,
- Gene expression profiles for each cell for each fish,
- Behavior traces,
- Stimulus time-series and information,
- Metadata consisting of information like cell masks.
- A second dataset is available also in figshare (https://figshare.com/s/72ceefe9844c1dda414a).
- Software, pipeline, and custom code generated for data processing are available on GitHub:

- Analysis code used to generate figures and associated information like gene names and clusters (https://github.com/zebrafish-WARP/WARP),
- Bigstream used for volumetric image registration (https://github.com/JaneliaSciComp/bigstream),
- Distributed Cellpose used for volumetric cell segmentation (part of https://github.com/MouseLand/cellpose, under cellpose.contrib.distributed_cellpose)
- Fishspot for detecting RNA transcript spots in volumetric images (https://github.com/GFleishman/fishspot),
- SpotDMix used for assigning RNA transcript spots to cells (https://github.com/Kepser/SpotDMix),
- segmentNMF for performing seeded demixed non-negative matrix factorization for extracting cell-resolution neuronal activity
- (https://github.com/L-Hess/segmentNMF),
- fish, a package for analyzing behavioral time series and calcium imaging data (https://github.com/d-v-b/fish).

## Supporting information

DataS1

## Acknowledgments

We gratefully acknowledge the shared resources and project teams at Howard Hughes Medical Institute (HHMI) Janelia Research Campus (JRC). We thank Janelia’s Project Technical Resources, led by Gudrun Ihrke, for support of EASI-FISH experiments (ME), as well as expert zebrafish dissections performed by Kari Close and Yisheng He. Two-photon and ExM light-sheet imaging experiments were conducted at the JRC Light Microscopy Facility. We would like to thank Damien Alcor and Michael DeSantis for microscopy training and assistance. We thank the Janelia Vivarium for animal care. We thank Haoze Zhu and Ilaria Gavetti for their manual annotations. We thank William E. Bishop for advice on data preprocessing and curation. We thank Scientific Computing, Stephan Preibisch, Janelia Experimental Technology, and the Janelia Visiting Scientist Program for support, and Ahrens lab members for discussions. We thank David Prober, Florian Engert, Vivek Jayaraman, Anoj Ilanges, Weiyu Chen, Gabriela Michel, and Javier How for comments on the manuscript. BE and LH received financial support through a VIDI grant to BE (016.Vidi.189.052).

## Author Contributions

Conceptualization: MA SS PT Data curation: EML ME GF LH Formal analysis: EML LH GF KS Funding acquisition: MA PT PK BE

Investigation: EML LH GF ME SN ABC Methodology: PT ME EML PK LH CS Project administration: MA

Resources: MA PK PT Software: EML GF LH KS Supervision: MA BE PT CS Validation: EML GF LH KS ME Visualization: EML GF LH

Writing – original draft: MA PT ME EML LH GF BE

Writing – review & editing: MA ME PT BM EML LH GF SS BE

## Declaration of Generative AI and AI-assisted technologies in the writing process

During the preparation of this work the author(s) used chatGPT in order to improve readability and language of the manuscript, and to improve code quality and readability. After using this tool/service, the author(s) reviewed and edited the content as needed and take(s) full responsibility for the content of the published article.

## Supplemental information

**Document S1** containing **Supplemental Figures S1-S12**.

**Data S1**. Spreadsheet with tabs containing: (i) List of EASI-FISH rounds, mRNA probes and HCR fluor; (ii) Spot count thresholds for each gene; (iii) List of subtypes for Figure 5B; (iv) Figure 5E Statistics; (v) Figure 5F Statistics; (vi) Figure 5G Statistics; (vii) List of subpopulations identified based on stimulus response, anatomy and subtype with their cell IDs; (viii) Statistics for LCD distributions for genes; (ix) Statistics for LCD distributions for clusters; (x) Figure 6H Statistics; (xi) Figure S12F Statistics.

**Video S1.** 3D representation of the experiment where a visual stimulus is presented to the fish. In this example we show the forward visual motion. Related to Figure 1.

**Video S2.** Four image volumes of the same fish brain showing 24 labeled genes. Related to Figure 1.

**Video S3.** Crops at different locations of the brain showing the genes probed in each of the 14 rounds. Related to Figure 1.

**Video S4.** 3D representation of the neuronal activity recorded (left) and the genes probed (right) in the same brain. Related to Figure 1.

**Video S5.** 3D representation of the neuronal activity recorded (changes in brightness) and the genes probed (color hue) in the same brain. In the top left corner we can see the *in situ* hybridization image in the expanded sample and below the ΔF/F of the time series max projection. Related to Figure 1.

**Video S6.** 3D representation of cell masks considered positive for *sst* in the thalamus. Related to Figure 2.

**Video S7.** Comparison of the raw time series (left) and the NMF reconstructed signals (right) and their ΔF/F. Related to Figure 2.

**Video S8.** Example subtypes for different genes and their average activity during visual stimulus presentations. Scale bar: 100 μm. Related to Figure 3.

**Video S9.** Subtypes involved in motor/swimming responses. Scale bar: 100 μm.

**Video S10.** Distribution of positive cells for each gene across the 3 fish. Scale bar: 100 μm. Related to Figure 4.

**Video S11.** The 15 largest subtypes that are positively or negatively correlated to the stimuli. Scale bar: 100 μm. Related to Figure 5.

## Methods

### Key resources table

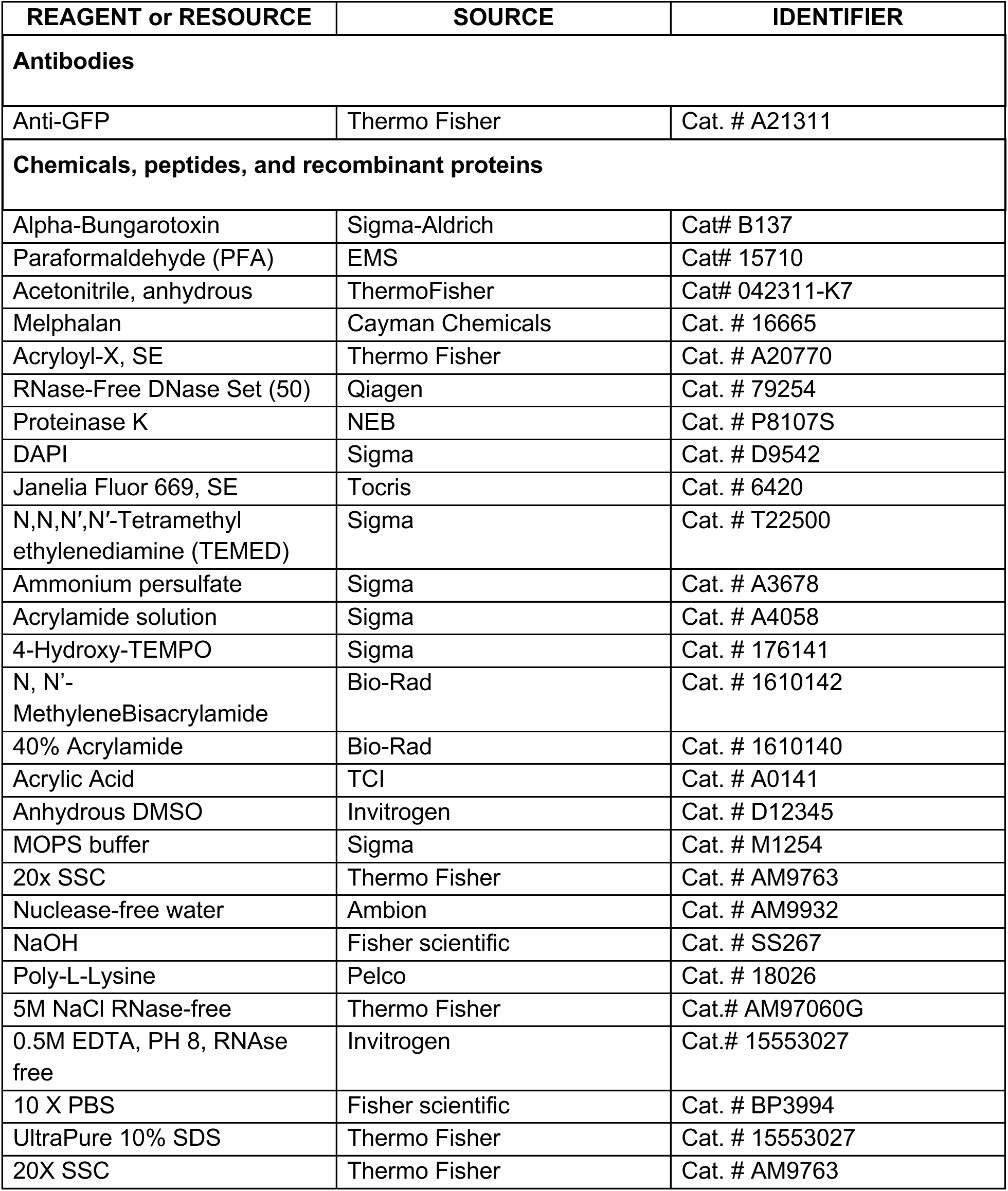
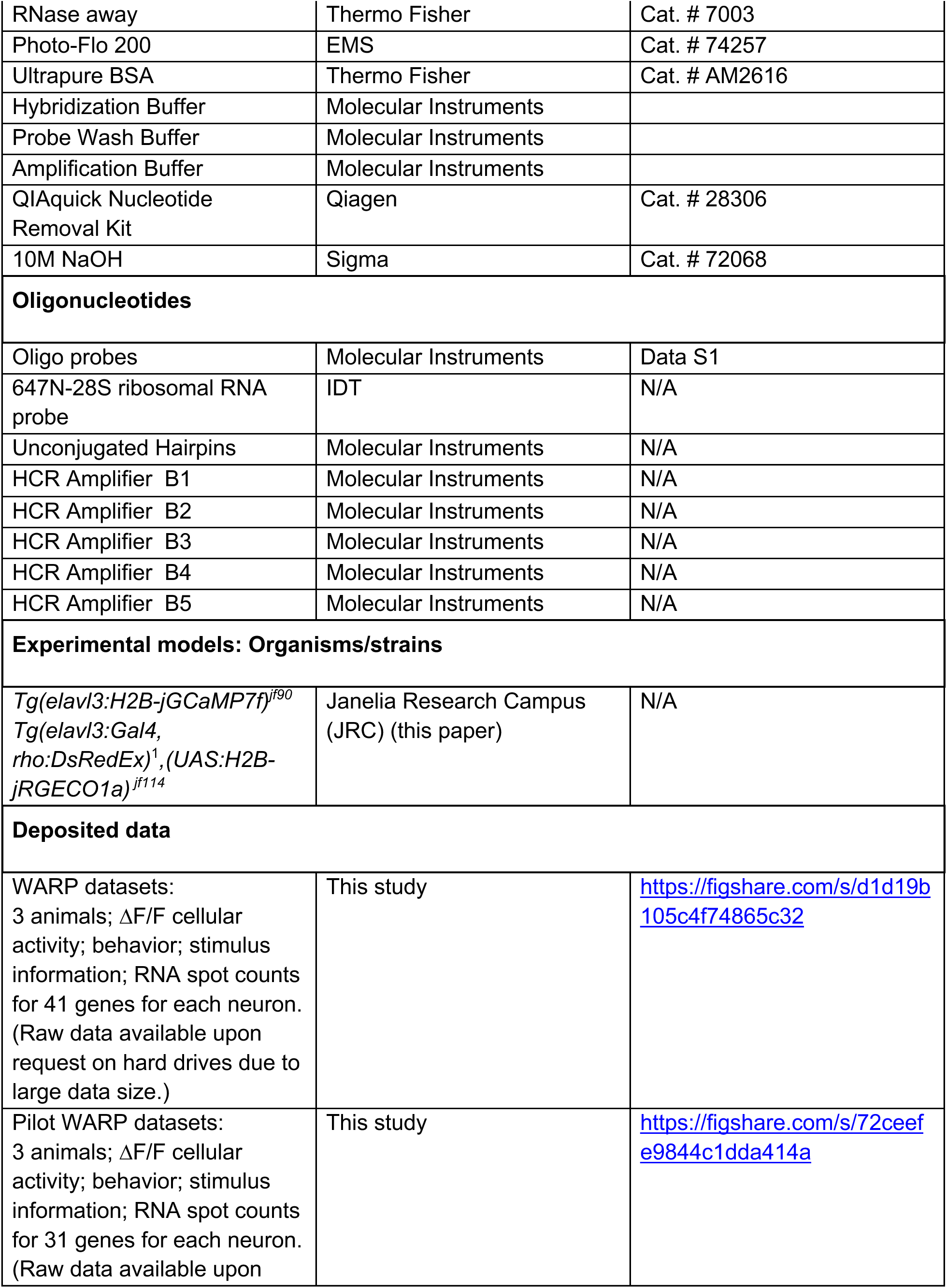
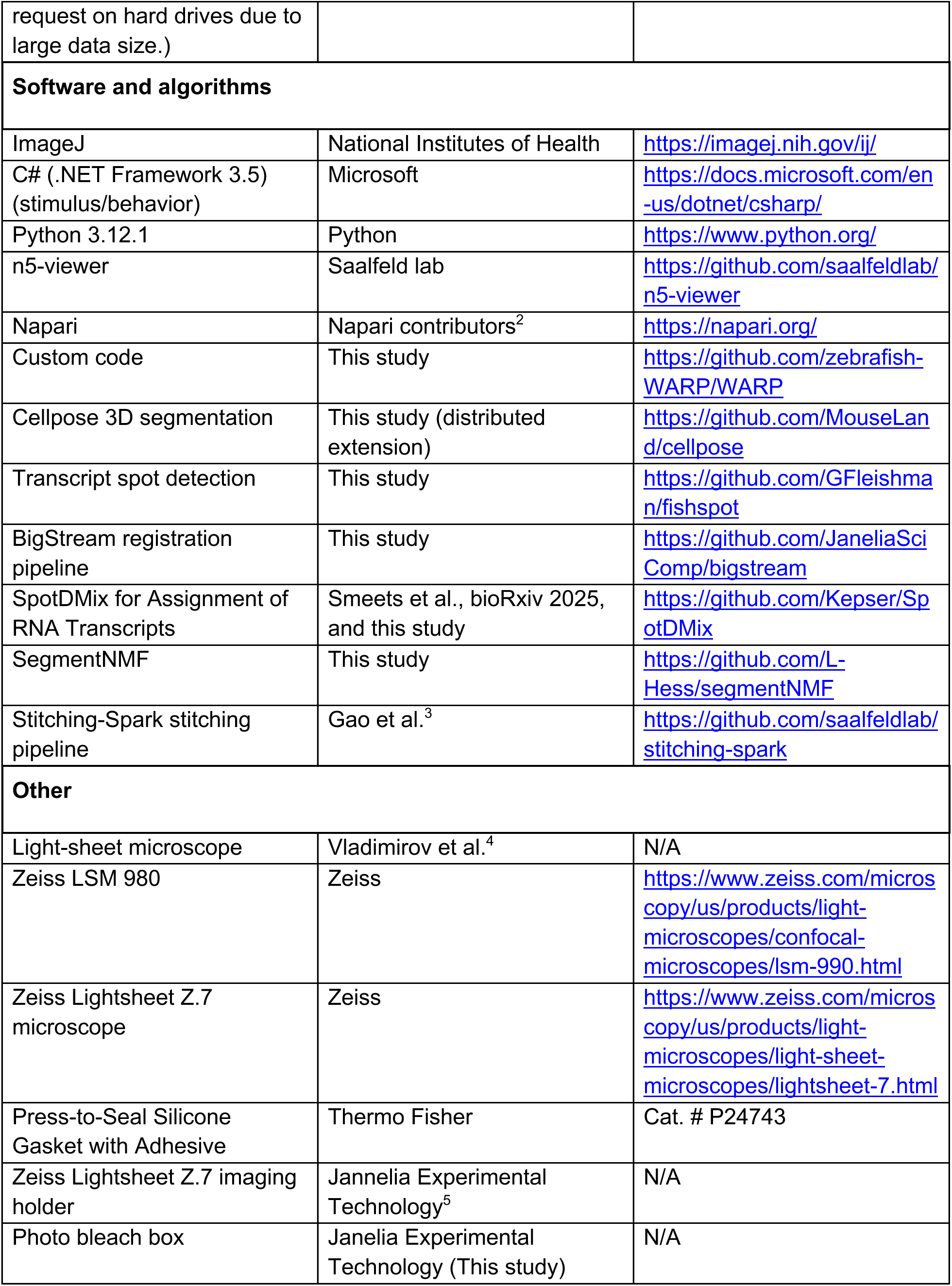

### Experimental procedures

#### Animals

Zebrafish larvae were reared at 28.5°C in 14:10 light-dark cycles. Animals were fed rotifers from 5dpf and used for experiments at 6dpf. At this age, sex cannot yet be determined.^6^ All experiments followed the protocols approved by the Institutional Animal Care and Use Committee at Janelia Research Campus.

The transgenic zebrafish larvae were in casper background^7^ and had a triple mutation after crossing *Tg(elavl3:H2B-GCaMP7f*) and *Tg(elavl3:Gal4, rho:DsRedEx),(UAS:H2B-jRGECO1a)* lines. These fish were expressing panneuronally the GCaMP7f^8^ calcium indicator in the nucleus under the elavl3 promoter. They were also expressing the red protein calcium indicator jRGECO1a^9^ in the nuclei of a random subpopulation of neurons using the gal4-UAS system and the elavl3 promoter.

#### Fictive behavior

As described previously,^4,10^ fish were paralyzed by 15 s immersion in 1 mg/mL in alpha-bungarotoxin (Sigma-Aldrich) and embedded with 2% low melting point agarose in a custom-build imaging chamber with an acrylic pedestal, glass walls and a diffusive screen at the bottom. Agar was cleared from part of the tail and two suction borosillicate pipettes (∼60 µm inner diameter, TW150-3, World Precision Instruments) were attached to the skin by gentle suction on each side of the tail. Extracellular motor nerve signals were amplified (Molecular Devices, Axon Multiclamp 700B) in current clamp and recorded with a custom written C# software. Signals were sampled at 6 kHz, and band-pass filtered with a 3 kHz/100 Hz low-pass/high-pass cutoff. “Swim vigor” was calculated as the standard deviation in a 10 ms sliding window. When swim vigor was above a 2.5 threshold, it was considered a “swim”.

#### Experiment sessions

Visual stimuli were presented to the fish using a custom-written C# software and projected (AKASO WT50 Mini Projector) to a diffuser at the bottom of the chamber. The projected light passed by a wratten filter (Red 25 Kodak Wratten 2) to be projected as red light. Experiments consisted of the following parts. 1. Recording of spontaneous activity for 10 min with no visual stimulus. 2. Positional homeostasis experiment^11^ for 10 min, in which a visual pattern drifted to produce OMR responses in closed-loop, followed by a pause and then a 1 s backward or forward drift was presented before returning to the closed-loop drift. 3. A train of visual stimuli repeated 10 times (23 min) where each stimulus was preceded by forward visual motion in closed-loop for 7 s, a pause for 2 s, a visual stimulus in open-loop for 3 s and a second pause for 5 s. The visual stimuli in open-loop were the following: forward visual motion, backward visual motion, rightward visual motion, leftward visual motion, light flash, dark flash, right loom (expanding dot) and a left loom. 4. A motor adaptation experiment^12^ where 10 s closed-loop visual motion at low-gain was alternated with visual motion at high-gain with 10 s pauses in between. 5. Open-loop visual motion presented for 10 min to produce swimming futility-induced passivity.^10^

#### Calcium imaging

Imaging of almost the entire brain was performed using a custom-build light-sheet microscope^4^. Briefly, two beams (front and side) were scanned horizontally and vertically to form laser sheets that scanned a volume of 350 µm in depth. A water-dipping 16x/0.8 NA objective (Nikon) was used to detect the emitted light which was recorded with a camera (Orca Flash 4.0 v2, Hamamatsu). At the beginning and end of each experiment, high-resolution stacks (∼100 ms exposure per plane) were taken at 1 µm steps in both green and red channels using 488 nm and 561 nm lasers and a band-pass filter (525/50 nm, Semrock) and long-pass filter (590 nm, Semrock) for separating GCaMP7f and jRGECO1a fluorescence. For the functional imaging only the frontal beam was used and 51 planes were acquired at ∼2 Hz with 7 µm steps using the 488 nm laser to record GCaMP7f fluorescence. At the end of each experiment, zebrafish larvae were fixed overnight in 4% PFA at 4°C.

#### Ex vivo two-photon anatomical reference

A volume of the whole brain was acquired with 2 µm intervals in z and 0.52 µm pixel size using a “Plan-Apochromat” 20x/1.0 DIC M27 75 mm water immersion objective (Zeiss) and two-photon imaging with the Zeiss LSM 980. The two-photon laser was tuned to 1010 nm; green and red channels were acquired using a non-descanned detector (NDD) with 505-545 and 575-605 nm emission filters, respectively. After this acquisition, the brain was dissected and stored in 70% ethanol at 4°C.

#### Selection of genes

The list of genes presented here for the 3 fish (fishID 59, 63, and 71, or Fish 1, 2, and 3 in main text) was selected based on an iterative optimization for gene expression located in the areas of active neurons, and then was extended based on the results of a first batch of experiments on which we targeted neuromodulatory genes. The final list was: *slc17a6b* (*vglut2a*), *gad1b*, *cx43*, *uts1*, *sst1.1 (*here named *sst)*, *pitx2*, *nr4a2a*, *pou4f2*, *cort* (*sst7*), *chata*, *cckb*, *tph2*, *gfra1a*, *calb2a*, *npy*, *crhb*, *dmbx1a*, *gbx2*, *pyya*, *nefma*, *chodl*, *pdyn*, *zic2a*, *th*, *esrrb*, *tbr1b*, *otpa*, *tac1*, *cart2*, *slc6a5* (*glyt2*), *grm1b*, *pvalb7*, *npb*, *slc6a2* (*net*), *irx1b*, *slc6a3* (*dat*), *penkb*, *calb1*, *penka*, *eomesa*, *emx3*, *cfos*.

With the aim to maximise the likelihood of target genes expressed in the neurons involved and responsive to our experimental assays, we first located the voxels of imaged neurons that correlated with swimming activity and responses to the visual stimuli. We used an NMF algorithm to segment cells based on their activity profiles,^10,11^ and then we calculated a Pearson correlation for each of the cells’ activity to vectors that indicated the timing of the stimulus presentations. The centroids of these ROIs were then registered to the mapZebrain atlas^13^ and a neuron size voxel (∼5 µm in diameter) was used for each of them to take an average of the fluorescence of 150 images of *in situ* hybridizations of the atlas.^14^ A sparse PCA with 30 components was then performed on the z-scored fluorescence of each of the ROIs and we identified the gene with the highest absolute loading for each of the components. This approach sometimes found genes expressed in non-neuronal populations or in very restricted locations. These genes were discarded from the original list and the process was repeated. After 5 iterations we picked the genes that repetitively appeared in the list of genes with the highest loadings of the PCA: *pitx2*, *uts1*, *nr4a2a*, *cort*, *pou4f2*, *chata*, *cckb*, *tph2*, *calb2a*, *npy*, *gfra1a*, *crhb*, *gbx2*, *dmbx1a*, *chodl*, *nefma*, *pyya*, *th*, *zic2a*, *pdyn*, *esrrb*, *otpa*, *tbr1b*, *glyt2*, *tac1*, *cart2*, *irx1b* and *npb*. To these genes we added the most common markers for glutamatergic and gabaergic neurons: *vglut2a*, *gad1b*. And *cx43* to locate glial astrocytes. Finally, based on our previous results and with the aim to locate neuromodulatory neurons and common anatomical markers we also included *sst1.1* (here named *sst*), *grm1b*, *pvalb7*, *calb2a*, *calb1*, *slc6a2* (*net*), *slc6a3* (*dat*), *penka*, *penkb*, *eomesa*, *emx3*. And we also included *fosab* (here named *cfos*) to locate strongly activated neurons. The labeling of *pitx2* was low and this gene was not processed for analysis.

The list of genes for of the first batch of experiments (fishID 50, 51, and 53, or Fish 4, 5, and 6) was selected based on a literature search and aiming to label neuromodulatory systems (neurons producing them and some of the receptors), locating non-neuronal populations and and including some common neuronal markers (e.g. calcium binding proteins): *slc17a6b* (*vglut2a*), *gad1b*, *cx43*, *sst1.1* (here named sst), *tph2*, *htr1aa*, *slc6a2*, *slc6a2* (*dat*), *sstr1a*, *npy1r*, *npy*, *slc6a9*, *glra1*, *chrna2a*, *chatb*, *grin1a*, *grm1b*, *pvalb7*, *gabra1*, *crhr1*, *crhb*, *calb1*, *calb2a*, *adra1bb*, *oprd1a*, *slc1a3b*, *aldh1a2*, *penkb*, *adra2b*, *penka*.

#### EASI-FISH

After being stored in 70% ethanol, the brains were embedded in a hydrogel according to the EASI-FISH protocol using a TREx-based recipe, except that we first omitted the initial DNAse digestion and acquired a “round zero” to register the ex vivo two-photon anatomical reference and EASI-FISH acquisitions. For round zero, hybridized with a directly conjugated 28S ribosomal probe (/5ATTO647NN/TG AGT TGT TAC ACA CTC CTT AG @ 1 µM) to label ribosomal RNA, stained all nuclei with DAPI and imaged residual GCaMP7f and jRGECO1a signal with a high (50%) laser power. The jRGECO1a signal for fishID 59, 63, and 71 (Fish 1, 2, and 3 in main text) bridged the ex vivo two-photon anatomical reference to round zero, the ribosomal probe signal bridged the ribosomal RNA signal in round zero to the total RNA signal in round one of EASI-FISH (stained with cyto-DAPI). The jRGECO1a signal ensured accurate registration across all three acquisitions (functional, two-photon and expanded). Before round one of EASI-FISH we DNAse digested the sample twice to digest genomic DNA and remove the ribosomal probe, and exposed the sample to a bright LED light (Cree LED LMH020-8000-35G9-00000TW) for 6hrs to photobleach jRGECO1a. EASI-FISH hybridization and Hybridization Chain Reaction (HCR) was performed as in Close et al.^5^, except that hairpins were used at 1:100. Gene probes used and respective lot numbers are listed in the Data S1. All fish were imaged in PBS (2x expansion) using multiple tiles on a Zeiss Z1 light sheet microscope.

Likely due to variations in expression levels and fixation details, we found that endogenous GCaMP7f signal and antigenicity (with Anti-GFP antibodies) stochastically varied from batch to batch. We were unable to see a usable GCaMP7f signal (endogenous or by antibody) for fishID 59, 63, and 71 (Fish 1, 2, and 3 in main text), but did for fishID 50, 51, and 53 (Fish 4, 5, and 6). While the GCaMP7f signal was not essential for registration, we used the CGaMP7f signal in fish 4-6 to validate our registration algorithms. For fish 4-6 the probe washes were shortened to half a day instead of overnight. With this shortened wash step, we observed a high level of spots in areas of the gel that did not have specimen, suggesting incomplete probe washing and potentially a higher level of background in the sample.

### Data preprocessing

#### EASI-FISH image stitching

The approximate dimensions of the 2× expanded brain are 600 x 2,000 x 1,000 µm. To cover this volume at an approximate sampling rate of 0.43 × 0.23 × 0.23 µm per voxel, EASI-FISH images are acquired as 5 × 3 mosaics of Z × 1920 × 1920 voxel tiles with an 8% spatial overlap between neighbors, where Z is large enough to cover the full dorsal-ventral axis. To accurately reconstruct a single continuous image, mosaics are stitched using the open-source stitching-spark software package (https://github.com/saalfeldlab/stitching-spark). Stitching-spark distributes the Fiji stitching plugin (https://github.com/fiji/Stitching) across a spark cluster so many large tiles can be aligned in parallel. Each tile is translated to match its neighbors, and a global optimization determines the best single translation for every tile. The optional flat field image intensity correction is not used, as for our particular data we found it added more artifacts than it removed. The stitched image is exported as a multi-scale pyramid in chunked N5 format with pre-computed down sampling factors along the (z, y, x) axes of: (1, 1, 1), (1, 2, 2), (2, 4, 4), (4, 8, 8), and so on. We refer to these scales as: scale 0, scale 1, scale 2, etc. We validated the accuracy of stitching qualitatively by inspecting overlap regions (Figure S1G) and quantitatively by computing the Pearon correlation of overlap regions (Figure S1H) before and after stitching.

#### Nucleus instance segmentation (distributed Cellpose)

Assignment of RNA transcripts to cells and extraction of neural activity from calcium images will utilize cell instance segmentation. The highest quality images of cellular anatomy available in our dataset are the total RNA and DAPI channels from EASI-FISH round 0 which label all soma and all nuclei respectively. Due to the high cell density of 6 d.p.f. zebrafish larva brain tissue, the boundaries between soma in the total RNA image are not sufficiently resolved to reliably perform cell body segmentation. However, the boundaries between nuclei at image sampling scale 0 are sufficiently resolved to perform nuclear instance segmentation, which proceeds in three stages: preprocessing, Cellpose segmentation, and postprocessing.

Preprocessing includes two steps: subtraction of the total RNA channel from the DAPI channel and unsharp filtering. The subtraction enhances contrast between nuclear boundaries by removing signal between them. Before subtraction, the total RNA channel intensities are uniformly scaled such that their Otsu threshold equals that of the DAPI channel. To further enhance nuclear boundaries, we apply an unsharp filter. Unsharp filtering is essentially high frequency amplification, and the filter width was chosen to amplify spatial frequencies whose wavelengths lie in the range of the typical nuclear diameter. This preprocessed image is segmented by the Cellpose 1.0 cyto2 model (note, not the nuclear model). We post-processed the segments for size. Any segment considered too small was merged with its largest surface area contact neighbor. Any segment considered too big was split with a watershed algorithm whose basin points were local minima of the distance transform to the segment boundary.

The scale 0 images used have approximate dimensions of 1,400 × 9,200 × 5,600 voxels. This is impractically large for the steps outlined above. Thus, the voxel grids were partitioned into overlapping blocks which were processed as described in parallel on distributed hardware (e.g. a cluster) and the results were smoothly stitched back together (Figure S1I). This distributed preprocessing and segmentation functionality is fully automated and provided by a single function which we have merged into the core Cellpose repository (https://github.com/MouseLand/cellpose).

We validated the instance segmentation quantitatively by comparing the number of segments found across six samples in 294 brain regions (Figure S1J). The Pearson correlation of segments per brain region between the samples is far higher than the correlation of segments per brain region to the brain region volume itself, indicating high consistency of nucleus instance detection.

#### EASI-FISH RNA transcript spot detection (FishSpot)

EASI-FISH rounds one and on each obtained a cyto-DAPI channel and three RNA transcript channels. Each labeled RNA transcript is assumed to be a point source of light smaller than the diffraction limit of the optics. Thus, we expect isolated transcripts to be imaged as point spread functions and clusters of transcripts to be imaged as superpositions of point spread functions with local maxima at transcript locations. RNA transcript detections are done with the full resolution scale 0 images. First, a white top-hat transform with a small structuring element is used to remove background signals. Second, the point spread function for the EASI-FISH round is estimated. This step must be repeated for every EASI-FISH round as the point spread function shape depends on the position of the light sheet relative to the objective focal plane, which can drift between acquisitions. To estimate the point spread function, a region dense with RNA transcripts is automatically detected. The local maxima in that region are identified and a small neighborhood around each maximum is extracted. The random sample consensus algorithm identifies a subset of these neighborhoods that average to produce a good representation of the point spread function. Third, to better separate clusters of RNA transcripts, this point spread function estimate is used to deconvolve the channel with the Richardson-Lucy algorithm. Finally, a Laplacian of Gaussian filter with a range of sizes assigns a unique 3D coordinate to the remaining local maxima. We validated spot detection quantitatively by comparing the number of spots detected across duplicated HCR rounds in 294 brain regions (Figure S1K). The Pearson correlation of detected spots per brain region between duplicated rounds is far higher than the correlation of detected spots per brain region to the brain region volume itself, indicating high sample and RNA integrity over time and high spot detection consistency.

### Assignment of RNA transcripts to nuclear segments (SpotDMix)

The combination of high density nuclear packing and the majority of RNA transcripts residing outside of the nucleus presents a unique challenge in assigning detected transcripts to nuclear segments. This prompted the development of a novel statistical method which models transcript locations as drawn from a weighted mixture model, in which each component in the mixture model is a distribution matching in shape to one of the nuclear segments.^15^

For this the shapes of the nuclear segments are approximated as ellipsoids obtained through a single-step fitting procedure per nuclear segment. Specifically, each ellipsoid is defined by an offset in space and a matrix defining a transformation of the unit sphere. These are obtained respectively as the center of mass and the Cholevsky decomposition of the covariance matrix of the voxel positions making up the nuclear segment. Finally, the resulting matrix is multiplied by an analytically derived factor to account for the uniform distribution of voxel positions inside the segment, up to voxel spacing per axis.

The mixture component of a nuclear segment is defined as a gamma distribution on the ‘ellipsoid distance’, i.e. a scaled version of the Mahalanobis distance with respect to the ellipsoid transformation matrix and offset, such that a distance of 1 corresponds to a point exactly on the ellipsoid surface. The gamma distribution is controlled by two global shape parameters modelling the peak location and distribution width respectively. Importantly, the shape parameters are shared globally across components to avoid local overfitting of the reach of single cells with no nearby transcripts.

All possible transcript-to-cell matches are included in a sparse distance matrix containing the ellipsoid distance from each transcript to each nuclear segment within a set distance threshold. Next, the mixture weights and the two shape parameters of the distribution are iteratively found through the Expectation Maximization (EM) algorithm. Membership probabilities are calculated after convergence and turned into ‘hard’ (as opposed to soft) assignments by assigning each transcript to the segment with largest membership probability.

The ellipsoid fitting procedure only needs to be performed once on the nuclear segments as a preprocessing step. After this, the EM algorithm can very efficiently be applied to the detected RNA transcript locations per imaged gene thanks to the round-to-round alignment. A more in-depth breakdown of the developed method is described in Smeets et al,^15^ including analytical results and a detailed performance evaluation on surrogate data sets.

### cF1 score and cAUPRC

In order to correctly estimate classification accuracy while dealing with the high number of negative cells, precision and recall were used, as they do not depend on the number of True Negatives after classification. We used a calibrated precision reference value of π_0 = 0.03 for all experiments, corresponding to a scenario where 3% of cells are positive.

The F_1 score measures a trade-off between precision and recall by taking the harmonic mean of the two values

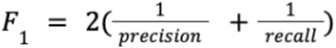

The calibrated F_1 score (cF1) is obtained by replacing the precision in this function with the calibrated precision, and is used throughout.

To better compared across cF_1 curves, we assigned a single scalar value to each method, a metric that measures the Area Under the Curve (AUC) in the (calibrated) Precision-Recall space (cAUPRC), where individual points correspond to a single threshold and the area summarises the performance for all thresholds.

### Bootstrapping and statistics of transcript assignment

Bootstrapping was done on the level of the output of the spot assignment algorithms by resampling with replacement the list of assigned spot counts, and comparing against the manual annotation labels with the same resampling.

For each gene the bootstrap distribution is obtained through n=100 resamplings with replacement. Error bars in figures show the standard deviation of this distribution.

The bootstrapped distributions for the cAUPRC score were used to perform statistical tests and calculate the effect size.

Comparing a method A to a method B is done through the pairwise distribution of resamplings; d(A, B) = mean(cAUPRC_A - cAUPRC_B)/std(cAUPRC_A - cAUPRC_B).

Reported p-values are Wilcoxon signed-rank test on the distribution of differences, alternative=greater, with Bonferroni correction on the number of comparisons (=18) (3 genes, 6 comparisons per gene)

### Manual annotation of transcript assignment

A small area in the thalamus was selected for manual annotation of cells (∼ 27 x 186 x 186 µm; 4075 cells), as genes like *sst*, *npy* and and *cx43* express close together. A positive/negative labelling was performed for three genes each by three annotators. The master annotation used in the analyses is obtained as the majority-vote between the three labels. Positive/negative classification was performed instead of individual spot assignment as this is often impossible to perform with certainty when there are neighbouring positive cells and is in general challenging for many spots.

Annotations were performed through a custom pipeline developed in Blender for easy navigation in three dimensions. Cells could be classified as positive, negative or unknown for the expression of one gene type. Classification was based on the raw fluorescent imaging data.

Inaccurate segments (e.g. too small) were omitted from the analysis. For the three genes, the number of omitted segments marked as unknown is (10, 3, 5) respectively.

### Naive transcript assignment methods

We used simple mask-based methods to assign spots. The first one is typically used in FISH methods and consists of only assigning spots that already overlap with the segmented cell, discarding spots located outside the mask of the cell. We called this method Segmentation Only in our comparative analysis. The second one is termed the Closest Centroid approach, where spots are assigned to the closest cell centroid. The third approach is termed Closest Mask where the spots are assigned to the closest mask. Spot locations were turned into indices for the volume containing the segmented masks. Around each spot, a box is extracted and distances are computed for all voxels which correspond to cells in this box. The spot is then assigned to the cell that has the minimum distance.

### Image registrations (BigStream)

The following sections cover all image registrations required to establish correspondence between EASI-FISH and calcium imaging. All registrations were solved by composing novel pipelines with BigStream (https://github.com/JaneliaSciComp/bigstream), an open-source python package for accurate registration of small features (e.g. cells) across huge images (e.g. tissues and whole samples). All images are foreground segmented prior to registration^16,17^; foreground masks ensure only voxels that are part of the sample affect registration. Optionally, any image with considerable blurring from scattered light can be unsharp filtered to enhance contrast at nuclear boundaries. Unsharp filtering is essentially high frequency amplification and kernel widths are chosen to amplify spatial wavelengths in the typical range of nuclear diameters. Finally, these image registrations are made more robust by the sparse jRGECO1a channel. For any alignment that is self-evidently correct the jRGECO1a channel can serve as an independent quantitative validation, but for any alignment that is not qualitatively satisfactory the jRGECO1a channel can be included in the registration itself to produce a more accurate result.

### EASI-FISH round N to round 0 registrations

The Round N cyto-DAPI channel is registered to the total RNA channel of round 0. The resultant transforms will be used to move RNA transcript detections to round 0 where they will be assigned to nuclear segments. Correct assignment cannot tolerate registration error larger than the typical distance between nuclear boundaries of 0.3 to 0.6 µm. Thus, we must achieve single voxel accurate registration at scale 1, which has an expansion corrected sampling rate of 0.21 × 0.23 × 0.23 µm. First, a global affine registration is performed at scale 3 to account for macroscopic misalignment due to physical sample handling. Second, we wish to perform local deformable registration of the scale 1 images which have approximate dimensions 1,400 × 4,600 × 2,800 voxels. This is impractically large for state-of-the-art deformable registration algorithms. Thus, the voxel grids are partitioned into overlapping blocks, affine and deformable registration is performed on each pair of blocks in parallel on distributed hardware (e.g. a cluster), and the resultant deformation fields are stitched into a single deformation of the entire scale 1 image. This distributed registration functionality is fully automated and provided by a single function call in the BigStream package. To validate registration performance, we manually annotated hundreds of cells for every registration performed. Using napari, a random crop from the fixed and registered image were presented and all cells in the crop were annotated as either correctly corresponding or incorrectly corresponding (Figure S1F). This was repeated for dozens of crops until more than 800 cells were annotated for each sample. Qualitatively, the fixed and registered images were superimposed and inspected for any areas where misalignment was visually apparent (Figure S1E).

### EASI-FISH round 0 to two-photon registration

The jRGECO1a channel of the round 0 EASI-FISH data is registered to the jRGECO1a channel of the two-photon data. Accuracy is limited by the two-photon z-plane/axial sampling rate of 2 µm, so we use the scale 3 EASI-FISH data, which has a sampling rate of 0.85 × 0.93 × 0.93 µm. This registration presents a unique challenge, as it must account for deformation introduced by the surgical removal of all tissue surrounding the brain and any deformation introduced by expansion microscopy preparation; both sample preparation steps must be done very carefully for this registration to have a chance to succeed. First, a global affine registration accounts for macroscopic misalignment due to physical sample handling. Second, the microsurgical procedure tends to introduce a low frequency bend and/or twist of the sample along its anterior/posterior axis. To correct such large but simple deformations, we divide the sample along the anterior-posterior axis into two overlapping blocks, affine register each separately, and smoothly stitch the result into a single deformation. This process is iteratively repeated with increasing numbers of blocks to gradually approach the smooth bend/twist. This nested piecewise affine algorithm is fully automated and available in BigStream as a single function. Finally, to correct high frequency deformation on the scale of individual cells, the entire image domain is broken into small overlapping blocks which are deformably aligned at full resolution. The results are smoothly stitched into a single deform of the entire image domain. To validate registration performance, we manually annotated over 1,000 cells for every sample. Using napari, a random crop from the fixed and registered image were presented and all cells in the crop were annotated as either correctly corresponding or incorrectly corresponding (Figure S1D). This was repeated for dozens of crops until more than 1,000 cells were annotated for each sample. Qualitatively, the fixed and registered images were superimposed and inspected for any areas where misalignment was visually apparent (Figure S1C).

### Two-photon to calcium imaging registration

The GCaMP7f channel of the two-photon data is registered to the GCaMP7f channel of the calcium imaging data. This registration presents a unique challenge, as it must account for deformation introduced by euthanasia and fixation and also register images with very different visual features due to the different microscopes used. For some samples, an unsharp filter was used to enhance contrast of individual cell boundaries in the calcium imaging reference image. First, a global affine registration accounts for macroscopic misalignment due to physical sample handling. Second, to correct high frequency deformation on the scale of individual cells, the entire image domain is broken into small overlapping blocks which are deformably aligned at full resolution. The results are smoothly stitched into a single deform of the entire image domain. To validate registration performance, we manually annotated over 1,000 cells for every sample. Using napari, a random crop from the fixed and registered image were presented and all cells in the crop were annotated as either correctly corresponding, incorrectly corresponding, or unclear (Figure S1B). Unclear cells are in areas where imaging quality is insufficient to determine if cells correctly or incorrectly correspond. This was repeated for dozens of crops until more than 1,000 cells were annotated for each sample. Qualitatively, the fixed and registered images were superimposed and inspected for any areas where misalignment was visually apparent (Figure S1A).

### EASI-FISH to calcium imaging registration refinement

Composition of the EASI-FISH round 0 to two-photon and two-photon to calcium imaging transforms puts EASI-FISH round 0 in register with the calcium imaging. However, those two datasets have not been directly compared. The composite transform is applied, and the resampled EASI-FISH GCaMP7f/jRGECO1a channel alignment is refined with a correction registration directly to the calcium imaging GCaMP7f/jRGECO1a channel. First, a global affine registration accounts for any residual macroscopic misalignment. Second, to refine alignment of individual cells, the entire image domain is broken into small overlapping blocks which are deformably aligned at full resolution. The results are smoothly stitched into a single deform of the entire image domain.

### Registration to the mapZebrain atlas

All registrations were solved using the tools and functions from BigStream (https://github.com/JaneliaSciComp/bigstream). We first made an “average brain” from our 6 samples. We registered all fish to a single fish and then registered them again to the average of the registrations in an iterative manner. The “average brain” was then registered to the mapZebrain atlas.^14^ Similarly to registrations described above, the GCaMP7f channel of the calcium imaging data images are foreground segmented prior to registration. Then, a global affine registration followed by a deform registration was performed. Finally, the transforms of the registrations were used to move the segments of the cells from their EASI-FISH round 0 coordinates to the atlas space and the centroids of these were located.

### Calcium imaging motion correction

Calcium imaging includes two image acquisition categories: functional time series volumes and anatomical reference volumes. Most of the imaging is the former which are parameterized for full brain coverage at two hertz; these volumes are low illumination and have a sampling rate of 7.0 × 0.406 × 0.406 µm. Anatomical reference volumes, which are collected once before functional time-series imaging and again after are higher illumination and have a sampling rate of 1.0 × 0.406 × 0.406 µm. We motion correct all time series data to the first anatomical reference. First, the anatomical reference is foreground masked. Second, the first frame of the time series data is affine registered to the anatomical reference. Third, all frames are affine registered to the first frame. Finally, all frames are resampled to the anatomical reference coordinate system. The resampled dataset is exported as a Zarr array with large time series chunks to facilitate downstream time series analysis.

### Segmentation initialized and regularized non-negative matrix factorization and ΔF/F calculation (SegmentNMF)

After image registrations the nuclear instance segmentation and calcium time series are aligned, which allows to take the contributions of each voxel assigned to a nucleus mask to obtain a single activity trace per cell. Following state-of-the-art calcium signal extraction we decided to use boundary spatial components that can overlap to model the possibility of signal mixing between neighboring cells.^18^ To this end, we developed a non-negative matrix factorization (NMF) algorithm that is initialized and regularized by the available anatomical segmentation.

Common implementations of NMF to extract cell signals from calcium data initialize the individual spatial components based on spatial footprints or spatial regions of high activity or correlation in the temporal domain.^18–20^ In our case, we initialize the spatial components directly from the aligned anatomical segmentation and the temporal components are null initialized, meaning no activity for any component at any time. The difference between the observed data and the model is optimized by alternating gradient descent steps along the directions of spatial and temporal components according to the Frobenius norm.

In order to de-mix signal from out of plane cells, we leverage the higher z resolution of the segmented volume (7 times more than the functional image). We applied a gaussian kernel with a given standard deviation (σ = 2 μm) to the higher resolution segmented volume. This procedure projects segments in-between planes onto the imaged planes and additionally increases the footprint of the spatial components in the x- and y-direction.

During optimization, we use a similar approach to regularize spatial components by weighting their gradients by distance from initialization with the same gaussian fall-off, i.e. spatial components change more slowly as they grow more distant from their initial shapes. At convergence, the spatial components have adapted to the time-series data and the temporal components have de-mixed activity of neighboring cells. The temporal components are retained for further study as the neural activity of each cell.

Due to the size of the time series, we partition the volume into overlapping blocks for which the NMF algorithm can be run in parallel on distributed hardware. Although possibly appearing in multiple blocks, each segment has only one ‘core’ block from which both spatial and temporal components are taken. All associated code for running the distributed segmentation initialized NMF can be found at https://github.com/L-Hess/segmentNMF.

To obtain the normalized change in fluorescence for each timestep (ΔF/F), we then take the resulting NMF temporal components and for each component divide it by a rolling baseline computed as the 10th percentile of the activity for that neuron in the preceding 120 time steps. In this process, we discard the first 120 frames for each experiment. These discarded timepoints coincide with the animal response and return to the baseline that follows the beginning of the laser scanning.

### Data analysis

For the cells to be considered positive for one of the genes. We visually inspected a binary classification of the spot counts with the signal from the HCR for each of the genes. We found that for most genes 20-25 was a suitable threshold of spots to get most of the cells that had clearly HCR signal around it while also avoiding false positive cells. Therefore, we used 25 as our minimum default threshold. For genes that had a wider distribution of spot counts (*’chata’*, *’glyt2’*, *’penka’*, *’cx43’*, *’cort’*, *’tph2’*, *’npy’*, *’gbx2’*, *’chodl’*, *’dat’*, *’npb’*, *’emx3’*, *’th’*, *’otpa’*, ‘*net’*), we used a Maximum Deviation approach to find the best suited threshold. Due to some sample and gene expression variability, some of those genes also had an adjusted threshold that was selected based on a careful comparison of the spot counts and the raw signal of the HCR. The thresholds for each gene and each fish (experiment) can be found in Data S1. For *cfos*, the threshold was 5 spots in order to be able to do quantitative comparisons.

### Molecular subtypes

To classify molecular subtypes, we constructed a binary matrix (neurons x genes) where cell positivity was determined by gene expression values exceeding a threshold. We primarily used a Maximum Deviation approach for threshold calculation, with manual adjustments made to account for variations in signal-to-noise ratio. (see above). We then only selected cells of each of the experiments (n=3) that were considered positive for at least one of the genes, excluding *cfos*. We then found the unique combinations of gene expression in these binary matrices (barcoding) and considered these as possible molecular subtypes. Each of these subtypes had to be composed of at least 5 cells that also have a minimum of 2 neighbors within a 40 µm radius. We further require each subtype to be found in each of the samples. Each subtype was assigned a unique color using distinctipy.^21^

### Anatomical and molecular subtype filtering

To ensure the replication of our results across the three animals, we conducted most analyses on a subset of neurons that met specific criteria. We selected cells from each molecular subtype across the three fish, excluding those positive for *cx43*. Within each of the 16 designated brain regions, we identified cells of the selected subtype, requiring a minimum of five cells from each fish to include that group for further analysis. Finally, we discarded any group in which a single fish contributed more than 70% of the total cells.

### Functional clusters

For each fish, we filtered the cells to include only those that were positive for at least one gene. This filtering focused on cells within the functionally acquired planes and included only those with activity detected after the segmented NMF step, excluding any traces that contained only NaNs or zeros. The selected cells were then clustered using Rastermap^22^ with the following parameters: n_clusters=30, n_PCs=50, locality=0.1, mean_time=False, applied to the session’s concatenated time series. Finally, for each of the 30 clusters, we applied the anatomical and molecular subtype filtering described earlier, retaining only cells of clusters that had genetic and anatomical replicates across all three fish.

### Plotting of matrices and vectors (Figure 3A)

To effectively handle the large number of neurons and enhance graphical representation, all vectors and matrices were binned every 10 cells (rows). For categorical variables—such as functional clusters, molecular subtypes, and brain regions—the representative row was determined by selecting the mode from the 10 corresponding cells. For quantitative variables, including activity traces, gene expression, and swimming correlation, the representative row was calculated as the average of the 10 corresponding cells. Furthermore, for the gene expression matrix, the first cell from each group of 45 or 130 was chosen to downsample the matrix according to the size of the matrix, ensuring it aligned in size with the other matrices.

### Swimming events (Figure 3A)

For each session of every fish, we calculated the windowed variance^4,10^ of the best of the two channels from the electrophysiological recordings of the tail. This measure is referred to as “swimming vigor.” We used swimming vigor to establish a threshold for detecting the start and end of swimming events. To classify these swimming events into categories of “swims,” “strong swims,” and “very strong swims,” we first identified the maximum value of swimming vigor within the window defined by the start and end of each swim. We then calculated the mean and standard deviation of these maximum swimming vigor values. Each swim was classified as follows: a “swim” if it was below the mean plus one standard deviation, a “strong swim” if it was above the mean plus one standard deviation, and a “very strong swim” if it exceeded the mean plus three standard deviations. Additionally, “no swims” were represented by the mean swimming vigor value recorded 100 time points (0.016 s) before each detected swim.

### Correlation to swimming behavior

To identify the cells that correlate with swimming activity, we downsampled the “swimming vigor” of each animal using a rolling average to align the time series with the imaging data (from 6 kHz to 2 Hz). We then calculated the Pearson correlation coefficient between these downsampled swimming traces and the activity of each neuron in their respective fish. This process yielded a correlation coefficient for each cell in each animal.

### Finding swimming related neurons

To select the swimming neurons at the bottom of Figure 3A, and for plots in Figure 3E-H, cells from the 3 fish that were positive for at least one gene and that had an absolute correlation coefficient higher than 0.1 were selected and then filtered with the anatomical and subtype steps mentioned above. For activity raster plots in Figure 3A and Figure S2B-C the cells of each fish were selected and ordered based on their functional cluster identity.

### Count of spots and positive cells across animals

For each fish, we identified the positive cells using the criteria mentioned above and calculated the mean and standard deviation of the spot count for the 41 genes across the entire brain (Figure S5A-B).

### Brain distribution of genes

For each fish we found the positive cells for each of the 40 genes (excluding *cfos*) in each of the 16 selected brain regions. If there were more than 10 cells, the number of positive cells was divided by the total number of cells in that brain region. These results are shown in Fig S5C. For Figure 4A we made an average of the 3 fish matrices.

### Co-expression matrices

For each of the fish, the positive cells for one gene (gene1) were selected and then, among those, we identified the positive cells for a second gene (gene2). We then calculated the ratio of the number of cells positive for both genes within the cells positive for gene1. We then calculated the average of across the 3 fish.

### t-SNE

To visualize the gene expression across the cells of the three samples, we performed a t-SNE^23^ in all the cells positive at least positive for one gene and their spot count for the 41 genes. We used the scikit-learn python package^24^ with the following parameters: n_components=2, verbose=1, perplexity=100, early_exaggeration=100, n_iter=1,000.

### Brain distribution of subtypes

We did the anatomical and subtype filtering as mentioned above (without filtering for *cx43* positive cells). For each molecular subtype that passed this filtering in each brain region, we calculated the mean number of cells across the three fish and divided it by the mean number of cells found in that brain region across the three fish.

### Spatial locality index and inner correlation for subtypes in brain regions

We did the anatomical and subtype filtering as mentioned above. Then the spatial locality of each of the identified molecular subtypes within each brain region was measured by means of a spatial locality index, adapted from Stringer et al.^25^ Specifically, we define the locality index as the Kullback–Leibler (KL) divergence between 1) discretized spatial distribution of neuron positions within the cluster, and 2) the discretized spatial distribution of all neurons positive for that gene across all animals. Neuron positions were binned into 3D spatial histograms using a 50 µm bin size along each axis (x, y, z), and histograms were normalized to form probability distributions. To normalize for the size of the different brain regions, we divided the spatial locality index by the number of discretized bins in that brain region (Figure S9A). The same group of cells was then divided by which fish they belonged to and a paired correlation matrix of the activity traces was calculated using Pearson correlation. Then, the lower triangle of the matrix below the diagonal was used to calculate the mean correlation of that group of neurons. These were finally averaged across fish (Figure S9B).

To find the subtypes that had high inner correlations and locality index we calculated the average of these measurements (average inner correlations = 0.2327, average locality index = 0.0021) across the brain and found the subtypes in each brain region whose values were higher (Figure S9C).

These paired measures of inner correlations and locality index per subtype were then analyzed with Pearson correlations brain-wide or by brain region (Figure S9D-F).

### Subtypes with 4 or more genes

Molecular subtypes with 4 or more genes were selected and followed by the anatomical and subtype filtering as mentioned above.

### Generating an activity averaged trial of the “Visual stimuli” session

During the Visual Stimuli session, the eight stimuli were presented sequentially in a stimulus train (as described above), repeated a total of ten times. We collected activity traces from the second to the tenth trials and averaged these nine trials to obtain a representative single trial for each neuron. We omitted the first trial as 120 timepoints were used for the calculation of the ΔF/F and therefore that trial was incomplete.

### Stimulus correlation analysis

The correlation between stimulus presentations and calcium activity traces from neurons was calculated using complementary approaches to account for variability in responses. First, we created a vector corresponding to the length of the visual stimuli time series, initially filled with zeros. We then placed ones in the vector from the onset of each stimulus presentation until two seconds after the stimulus disappeared. For each stimulus, we calculated the Pearson correlation between its respective vector and the activity trace of each neuron obtained from the segmented non-negative matrix factorization (NMF). Neurons that were not in the functional imaging planes or that showed no signal after the NMF method were excluded from this analysis.

To address the possibility that some neurons might respond to the stimulus with a delayed reaction rather than at the exact moment of stimulus onset (for example, in cases of swimming responses or longer perception times for looming stimuli), we created another set of vectors similar to the first, but with the onset delayed by two seconds. We calculated Pearson correlations for these delayed vectors in the same manner as for the initial set. The results from both correlation analyses were combined by retaining the maximum absolute value from each.

This initial approach is particularly effective for identifying cells that are specifically tuned to individual stimuli, as activity occurring between presentations can reduce the correlation coefficient. To also consider neurons that might respond to multiple stimuli (e.g., both a dark flash and a loom, which share a dimming component) or to some but not all stimulus presentations, we conducted Pearson correlation analyses over 10-second windows, with three seconds before and seven seconds after each individual stimulus presentation. This windowed analysis was also performed using the delayed vectors. We then calculated the median correlation coefficient for each stimulus across its presentations. Similar to the two sets of correlation analyses with the full vectors, the windowed results were combined by taking the maximum absolute value between the regular and delayed approaches.

### Selecting positively and negative correlated neurons

To identify cells that responded to the stimuli, we first calculated the coefficients at the 90th percentile for each stimulus and then determined their average for both the full stimulus vector (0.129) and the windowed results (0.434). When filtering for positively correlated cells, we selected those with a correlation coefficient greater than the respective means from either the full vector or the windowed results. For negatively correlated cells, we chose those with a correlation coefficient lower than the negative of these means (−0.129 and −0.434). When selecting for both positively and negatively correlated cells, we included cells with a correlation coefficient that exceeded the maximum absolute value of the means.

After this first selection, cells would be filtered by the anatomical and subtype filtering as mentioned above. For Figures 5B,C and D the molecular subtypes with just one gene were also filtered out.

### Plotting location, activity and behavioral responses of forward visual motion related subtypes

The selection of the 15 largest molecular subtypes, which had the highest number of responsive cells to the forward visual motion stimulus, was performed as previously described. The 3D coordinates of these neurons in atlas space were then plotted, and their averaged activity trace for each molecular subtype was calculated from the representative trial mentioned earlier. This trace was plotted within a window that included five seconds of the closed-loop period, followed by a two-second pause, the stimulus presentation, and three seconds after the stimulus presentation.

To visualize the behavioral responses, we calculated the “swimming vigor” for each fish, as described earlier. The trials for the forward visual motion (from the second to the tenth) were then overlaid and aligned with the timings of the calcium activity.

### gylt2_gra1a analysis

After the analysis that found their presence (Figure 5A-B), we selectively searched for the gylt2_gra1a neurons in the inferior medulla oblongata that had higher correlation coefficients than 0.129 to the full stimulus vector. Their location in atlas space was then plotted. To compare their responses to the ones from other cells within the medulla oblongata, we also selected the remaining responding cells and the non-responding cells from that region. The average of their maximum ΔF/F value between the 3 seconds prior and the 7 seconds after the onset of the stimulus was compared across these three groups with a Kruskal-Wallis H-test. If the test was significantly positive, a post hoc pairwise test for multiple comparisons of mean rank sums (Dunn’s test) was performed with a Bonferroni correction. The summary of these tests are indicated in Figure 5E. The same analysis was performed for their correlation to swimming vigor. For alternating trials, we plotted the average activity of the selected gylt2_gra1a over the “swim vigor” of a fish.

### pou4f2_cckb analysis

After the analysis that found their presence (Figure 5 A-B), we selectively searched for the pou4f2_cckb neurons in the periventricular area that had higher correlation coefficients than 0.129 to the full stimulus vector. Their location in atlas space and their average activity trace from the averaged trail (see above) was then plotted. To compare their responses to the other stimuli, the average of their maximum ΔF/F value between the 3 seconds prior and the 7 seconds after the onset of the stimulus was compared across stimuli with a Kruskal-Wallis H-test. If the test was significantly positive, a post hoc pairwise test for multiple comparisons of mean rank sums (Dunn’s test) was performed with a Bonferroni correction. The summary of these tests are indicated in Figure 5F.

### Subtypes correlation to stimuli matrix

For each stimulus, we identified responsive cells by selecting both positively and negatively correlated cells. This selection included cells with a correlation coefficient that exceeded the maximum absolute values of the means for both the full stimulus vector (0.129) and the windowed results (0.434). We then applied anatomical and subtype filtering as previously described. If a sufficient number of cells met these criteria (>=5), we calculated the average correlation coefficient for the neurons of each fish within each molecular subtype and brain region, and subsequently averaged these values across all fish. In this way we could define neuronal subpopulations based on their responses to the stimuli, their subtype and the anatomical region they are located in.

### *cfos* labeling and analysis

After the experiment was completed, the fish were euthanized and fixed in 4% paraformaldehyde (PFA). The fixation was planned to occur approximately 30 minutes after the end of the Visual Stimuli session, as labeling neuronal activity by detecting *cfos* mRNA is most effective around this time following sensory stimulation or behavior.^14,26^ We defined cells as *cfos*-if they had fewer than 5 mRNA spots assigned to them, and as *cfos*+ if they had 25 spots or more.

To compare their maximum response (ΔF/F), standard deviation, maximum correlation to stimuli (windowed correlations), and correlation to swimming, we used the Mann-Whitney U test. We also employed the Mann-Whitney U test to compare the windowed correlation coefficients of negative and positive *cfos* cells for each stimulus presentation, applying a Bonferroni correction of 0.05/8 = 0.00625.

To assess the number of *cfos*+ cells per brain region, their proportion within molecular subtypes, and the number of spots, we first selected cells with 25 or more spots. We then performed anatomical and subtype filtering as described above, with the exception that each subtype needed to have more than one cell (instead of five) per fish to be included. Finally, these measures were averaged across fish.

### Calculation of Local Correlation Difference

We quantified how strongly neurons sharing a gene are coordinated relative to their local neighborhood using the Local Correlation Difference (LCD).

For each fish and gene *g*, we first identified the set *S*_g_ of neurons that express *g* (for the binarization method, see Methods: Identifying gene-positive cells). Let *y*_i_(*t*) denote the calcium activity of neuron *i* over time. We computed a Pearson correlation matrix between all neurons in *S*_g_ and all neurons in the brain. For each pair of gene-expressing neurons (*i*, *j*) with *i*, *j* ∈ *S*_g_, we treated neuron *j* as the spatial “anchor” and defined:

- *C_ij_*: the Pearson correlation between the activity of neurons *i* and *j*;
- 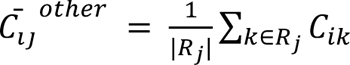: the average correlation between neuron *i* and the non-gene-expressing neighbors of *j*. Here *R*_j_ denotes the set of spatial neighbors of neuron *j* that lie within 15 µm and do not express gene *g*.

We then define Local Correlation Difference for the pair (*i*, *j*) as:

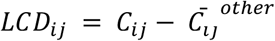

Thus, for each anchor neuron *j*, *LCD*_ij_ measures how strongly a gene-matched partner *i*correlates with *j* compared to how strongly *i* correlates with nearby neurons around *j* that do not express the gene.

LCD values were computed for all ordered pairs of neurons in *S*_g_ (excluding same neuron pairs (*i*, *i*), which were set to NaN), yielding an *N*_g_ × *N*_g_ matrix per gene and fish with *N*_g_ denoting the number of gene-positive neurons for that fish. To avoid noisy or unstable estimates, we excluded anchors *j* with fewer than 5 neighbors in *R*_j_; all pairs (*i*, *j*) with such anchors were also set to NaN. A neighborhood of 15 µm was chosen to balance capturing local correlations while ensuring most neurons had sufficient neighbors for stable calculation of the metric.

As individual pairwise LCD values proved noisy and to obtain a per-neuron statistic, we summarized the LCD at the level of single neurons and spatial bins. For each neuron, we computed a distance-restricted average LCD by averaging *LCD*_ij_ over gene-matched partners *j* ∈ *S*_g_ within a spherical neighborhood of radius 20 µm, subject to a minimum of 5 partners in the bin. (Figure S12A). These per-neuron, distance-binned LCD values, *LCD*_i_(*t*) were used for downstream analyses and denote LCD.

Further to note, all analyses were performed at the level of gene-types (gene-positive vs gene-negative), rather than finer molecular subtypes. This choice was motivated by the fact that many molecularly defined subtypes were too sparse locally to yield stable neighborhood averages.

### Permutation-based null model for Local Correlation Differences

To test whether observed LCD values reflect gene-specific structure beyond what would be expected from local anatomy and generic correlation structure, we constructed a permutation-based null model for each gene and fish.

The key idea is to locally shuffle gene identity within spatial neighborhoods while preserving the number and spatial density of gene-positive neurons and the empirical activity correlations. Concretely, for each gene *g* and fish:

1. For each gene-expressing neuron *j* ∈ *S*_g_, we computed:

its set of non-gene-expressing neighbors *R*_j_ within 15 µm, and
the corresponding LCD matrix *LCD*_ij_ as defined above.
2. For each permutation, we created a “pseudo-gene” configuration by reassigning the spatial anchor positions of gene-expressing neurons:

For every gene-expressing neuron *j* ∈ *S*_g_, we randomly selected a neuron *j*^(1)^ from its local neighborhood (within 15 µm), and treated *j*^(1)^ as the pseudo gene-matched anchor for that permutation.
For each such pseudo anchor *j*^(1)^, we then sampled a set of local neighbors *R*_#_(#) from its own spatial neighborhood with replacement, choosing the same number of neighbors as in the observed configuration (|*R*_j_(#) | = |*R*_j_|). Importantly, the choice of pseudo anchors and neighbors was irrespective of positivity for genes and the underlying activity time series and the anatomical positions of all neurons were left unchanged.
3. Using exactly the same formula and neighborhood radius as for the observed data, we recomputed

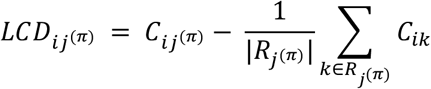

for all *i* ∈ *S*_g_ and all pseudo anchors *j*^(1)^. This yields an LCD matrix for each permutation that respects the spatial organization and local correlation statistics of the data but breaks the alignment between gene expression and correlation structure.

We repeated this procedure for 2,500 permutations per gene and fish, generating an empirical null distribution. For each gene, we then computed the same neighbor-averaged statistic *LCD*_ij_ from the observed LCD matrices and from each permuted LCD matrix. Empirical two-sided p-values were obtained by comparing observed statistics to their permutation distributions within each fish; Benjamini-Hochberg correction was applied to the p-values across genes. As p-values were calculated for individual fish, we took the largest value across fish as a conservative estimate of the full significance and reported those.

### Spatial clustering of Local Correlation Differences

To identify spatially localized neuronal clusters associated with gene-specific LCD patterns, we clustered the spatial distribution of gene-expressing neurons and then summarized LCD values within each spatial cluster.

For each gene, we first pooled all neurons that positively expressed that gene across all fish and extracted their 3D locations in MapZebrain-aligned coordinates. To avoid splitting bilaterally symmetric clusters across hemispheres, we projected neurons from the right hemisphere onto the left prior to constructing spatial neighborhoods. Specifically, we mirrored the lateral (left–right) coordinate of right-hemisphere neurons across the midline of the brain estimated from the neurons in all fish using a symmetric projection.

We then constructed a k-nearest-neighbor (kNN) graph on these projected coordinates using the kneighbors_graph function (k = 25) in Scikit-Learn^24^. On this graph, we ran the Leiden community detection^27^ to obtain compact spatial clusters. The effective granularity of the clusters was controlled by a target size of approximately 10 neurons per cluster and a resolution parameter of 0.1.

We then quantified the contribution in numbers of neurons of each fish to each cluster and defined a fish-support criterion to ensure clusters reflected gene–function relationships reproducible across animals. A cluster was marked as “fish-supported” if (i) it contained at least 25 neurons (≥ 25), and (ii) each fish contributed at least a 15% of the neurons relative to the uniform share. In downstream analyses we focused only on clusters that passed this fish-support criterion.

For each cluster, we aggregated the LCD values of its member neurons to obtain a cluster-level median LCD. For each cluster, we further aggregated the permuted LCD values over the same neurons and calculated p-values in the same manner as for the full gene distributions; again Benjamini-Hochberg correction was applied, here across clusters.

### Time-resolved LCD contributions

To obtain time-resolved LCD contributions, we note that, since the LCD consists of a difference in Pearson correlations, it is possible to decompose the contributions to the total metric in time. For each fish and gene *g*, let *S*_!_ be the set of gene-positive neurons and *y*_i_(*t*) the calcium activity of neuron *i* at time *t*. Pearson correlations can be written as a temporal average of z-scored co-fluctuations:

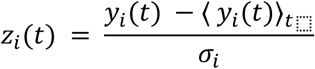

Then, for any pair of neurons (*i*, *j*):

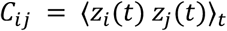

Using the same anchor-neighborhood definition as in the static LCD (anchor *j* ∈ *S*_i_, non-gene neighbors *R*_j_ within 15 μm), the *other* correlation term can be rewritten as:

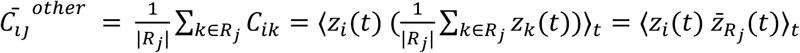

We can therefore define per-timepoint correlation contributions as:

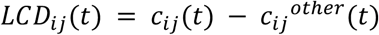

and the per-timepoint LCD contribution as:

Again, to obtain stable estimates, we averaged *LCD*_ij_(*t*) (or separately *c*_ij_(*t*) and *c*_ij_^other^(*t*)) over the same set of pairs (*i, j*) over neighbors *j* within a 20 μm radius to obtain *LCD*_ij_(*t*), and report the average per-timepoint LCD contributions across stimulus presentations for the neurons in each cluster.

To obtain correlations with swimming intent, we calculated the Pearson correlation between the neighbor-averaged per-time-point LCD contribution of all neurons in a cluster and the swimming intent downsampled to match the calcium time series (calcium activity and behavior matched per fish). Downsampling was performed by taking the maximum power of the electrophysiological signal in each time bin.

## WARP links 2025

Figshare datasets:

Main dataset: **Error! Hyperlink reference not valid.** Pilot dataset: https://figshare.com/s/72ceefe9844c1dda414a

GitHub Custom code:

Analysis: https://github.com/zebrafish-WARP/WARP

Cellpose 3D segmentation: https://github.com/MouseLand/cellpose Transcript spot detection: https://github.com/GFleishman/fishspot

BigStream registration pipeline: https://github.com/JaneliaSciComp/bigstream SpotDMix for Assignment of RNA Transcripts: https://github.com/Kepser/SpotDMix SegmentNMF: https://github.com/L-Hess/segmentNMF

## Supplemental Figures

**Figure S1:**
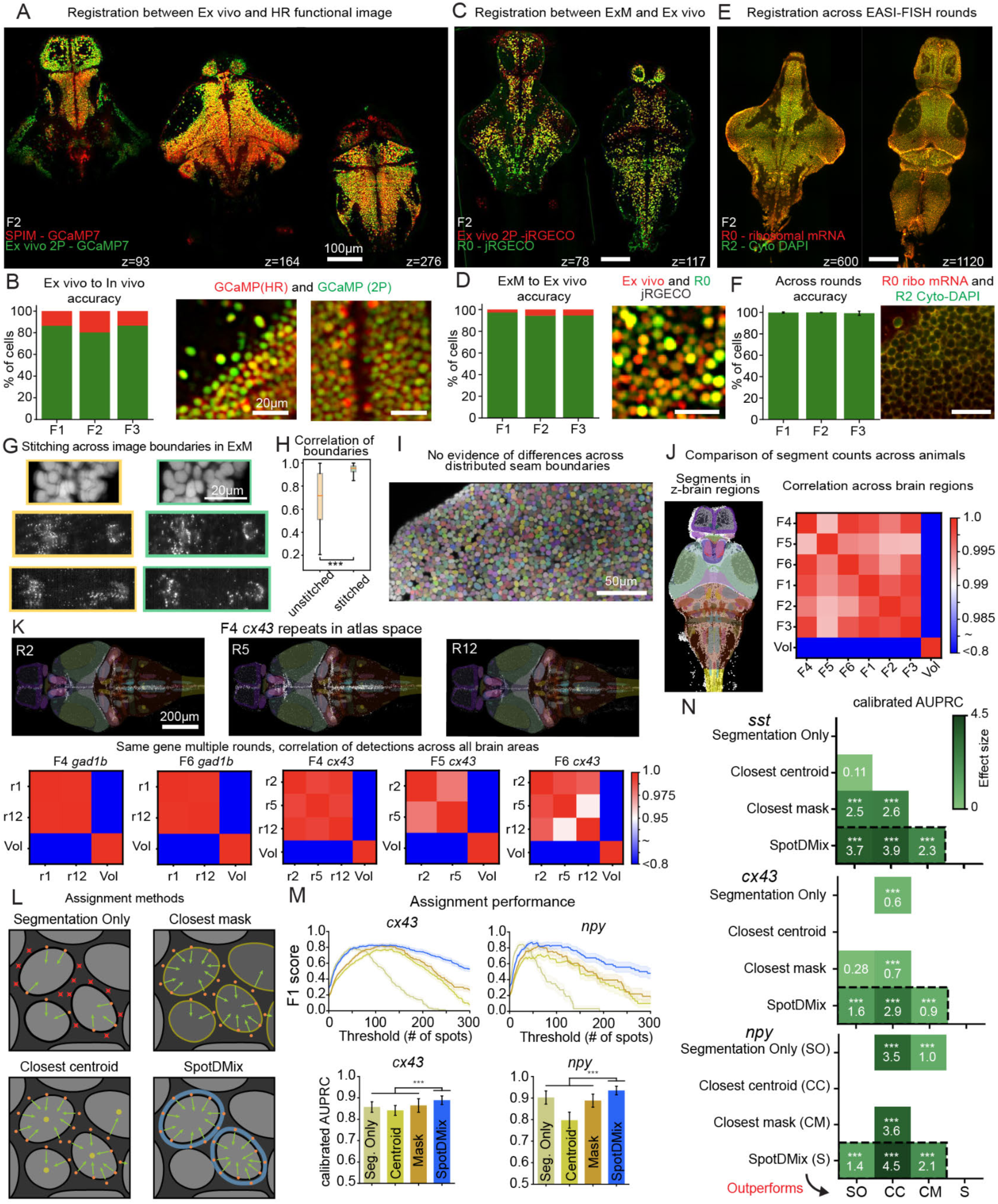
Computational tools (A) Three planes showing overlapping images after registering the ex vivo two-photon volume (green) of Fish2 to the anatomical reference of the in vivo (red). (B) Left: Proportion of accurately and incorrectly registered cells for the 3 fish. This proportion is taken from the cells that were either clearly correct or clearly incorrectly registered (Total number of annotated cells: F1 = 1176, F2 = 1060, F3 = 1025; Proportion of clear cells: F1 = 75.17%, F2 = 80.18%, F3 = 88.78%). Right: Zoomed-in crops of the registered images for F2. (C) Two planes showing overlapping images after registering the expanded sample (green) of Fish2 to the anatomical reference of the Ex vivo (red) for the jRGECO labeled cells. (D) Left: Proportion of accurately and incorrectly registered cells for the 3fish. This proportion is taken from the cells that were either clearly correct or clearly incorrectly registered. (Total number of annotated cells: F1 = 1022, F2 = 1070, F3 = 1029; Proportion of clear cells: F1 = 96.87%, F2 = 97.75%, F3 = 95.72%). Right: Zoomed-in crop of the registered images for F2. (E) Two planes showing overlapping images after registering the expanded sample after round 2 (green) of Fish2 to the expanded sample after round 0 (red) for mRNA staining with probes or DAPI. (F) Left: Average proportion of accurately registered cells for the 3 fish across the 14 rounds. This proportion is taken from the cells that were either clearly correct or clearly incorrectly registered (Average and standard deviation of number of annotated cells per round: F1 = 1054.35 (367.24); F2 = 856.64 (340.76); F3 = 884.71 (249.55); Average and standard deviation of % of clear cells per round: F1 = 99.88 (0.31); F2 = 99.88 (0.25); F3 = 100 (5.37e-15)). Right: Zoomed-in crop of the registered images for F2. (G) Example image crops of expanded sample before (left) and after (right) stitching. (H) Correlation of overlapping regions before and after stitching. (Wilcoxon signed-rank test; ns p>0.05; * p<0.05; ** p<0.01; *** p<0.001). (I) Crop showing segmented cells after using distributed cellpose. Note that no linear boundaries separating segments are observed indicating that the stitching of the segments was successful. (J) Left: Brain regions of the atlas over the detected segments of one of the samples. Right: Comparison of number of segments found within anatomically defined brain regions across animals. A Pearson correlation comparing each fish and the volume of the brain areas (vol). Note that all the fish have a correlation coefficient of 0.99 or higher, and that the correlation to the volume of brain areas is smaller than the one across fish. (K) Comparison of detected mRNA transcripts across multiple rounds performed in another dataset (Fish 4, 5and 6). Top: Detected transcripts for cx43 in rounds 2, 5 and 12. The brain regions of the atlas are overlaid. Bottom: Comparison of number of transcript detections found within anatomically defined brain regions across rounds. A Pearson correlation comparing each fish and the volume of the brain areas (vol). Note that all the fish have a correlation coefficient of 0.95 or higher, and that the correlation to the volume of brain areas is smaller than the one across fish. (L) Schematic representation of the naive methods and SpotDMix. (M) Top: curves of F1 score for different thresholds of number of spots for cx43 and npy. Bottom: calibrated AUPRC for cx43 and npy. SpotDMix outperforms the other methods significantly (Wilcoxon signed-rank test with Bonferroni correction; ns p>0.05; * p<0.05; ** p<0.01; *** p<0.001). (N) Calibrated AUPRC paired comparisons of the mRNA transcript assignment methods. Only comparisons where the rows outperform the columns are shown (Wilcoxon signed-rank test with Bonferroni correction; ns p>0.05; * p<0.05; ** p<0.01; *** p<0.001).

**Figure S2:**
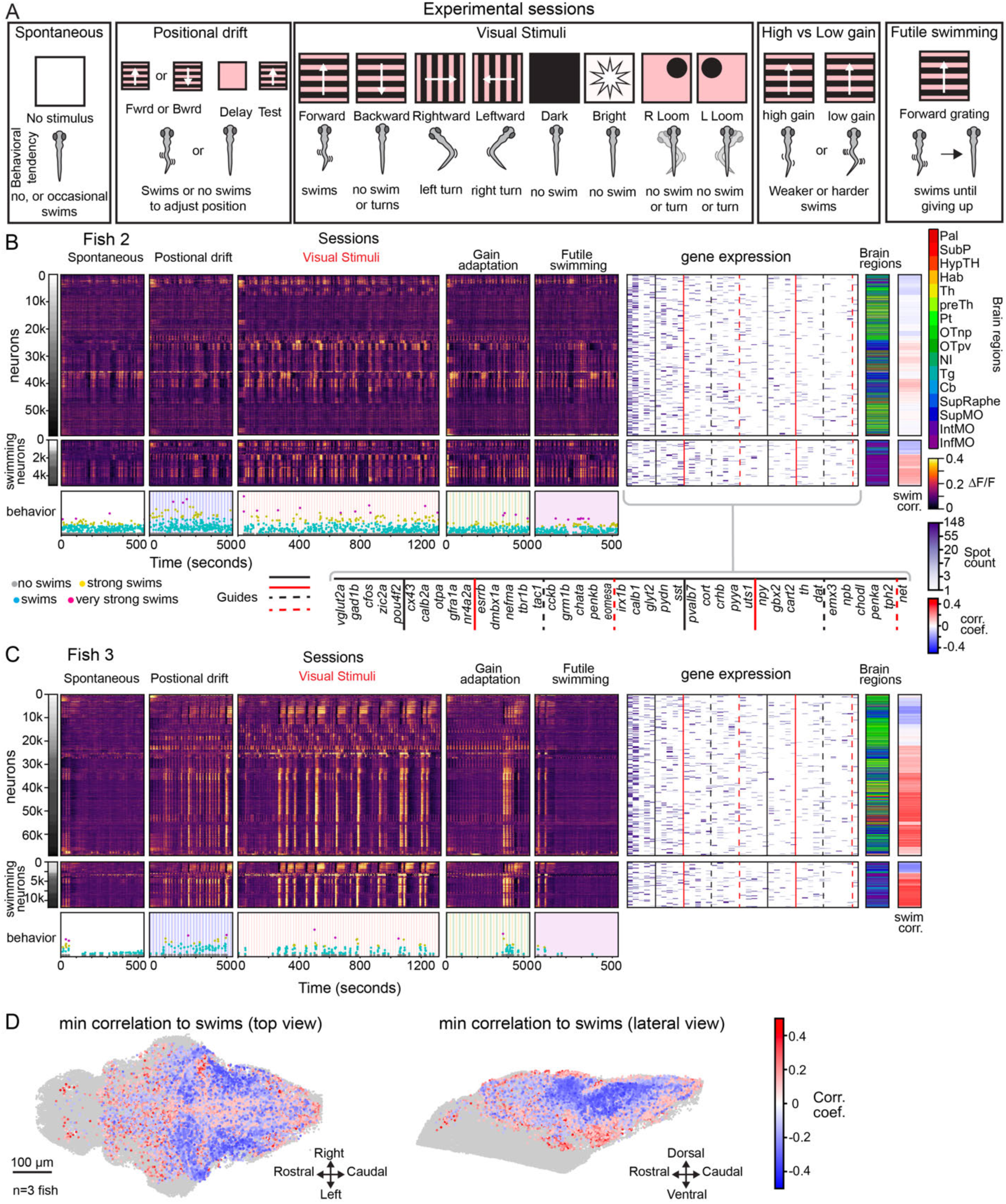
Multimodal datasets (A) Schematic representation of the experimental sessions, the visual stimuli included and the behavioral responses that they tend to produce. For details see Methods: Experiment sessions. (B) Overview of the dataset for an example fish (Fish 2). The top row of matrices and vectors are representing the different features, ordered by their similarity of activity during all sessions. The first vector represents the functional cluster (k=30). The next 5 matrices represent the neuronal activity traces (ΔF/F) for each of the sessions of the experiment. The following matrix represents the spot counts of the 41 genes. The Brain region vector represents the location of the neurons based in a brain atlas. The last vector represents the correlation values of the neuronal activity with the swimming behavior (Methods: Correlation to swimming behavior). All vectors and matrices were binned every 10 cells (rows) for clarity (See Methods: Plotting of matrices and vectors (Figure 3A)). The middle row of matrices and vectors show the same as above, but for cells selected from their correlation to the swimming activity (R>0.1 and <-0.1). The bottom row shows the swimming responses for each of the sessions of the experiment. (C) Overview of the dataset for an example fish (Fish 3). The organization of matrices and rows is the same as in B. (D) Correlation coefficient values of the neuron activity traces and the swimming activity. Ordered to plot the lower values on top (minimum projection).

**Figure S3:**
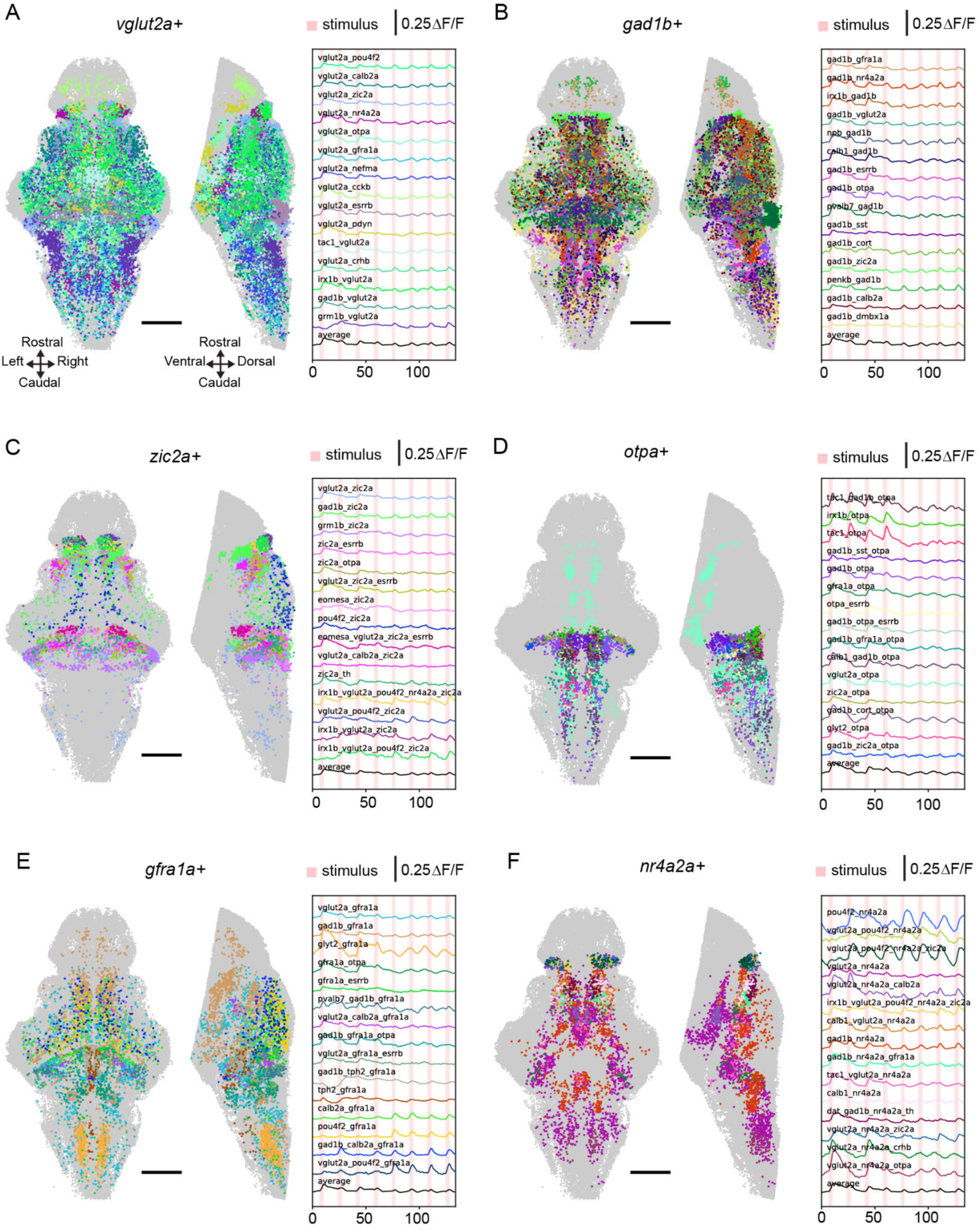
Example molecular subtypes (A-F). Brain maps showing the cells from the 15 largest molecular subtypes with expression of one main gene and another gene. The right panel shows the average activity of the neurons in these subtypes.

**Figure S4:**
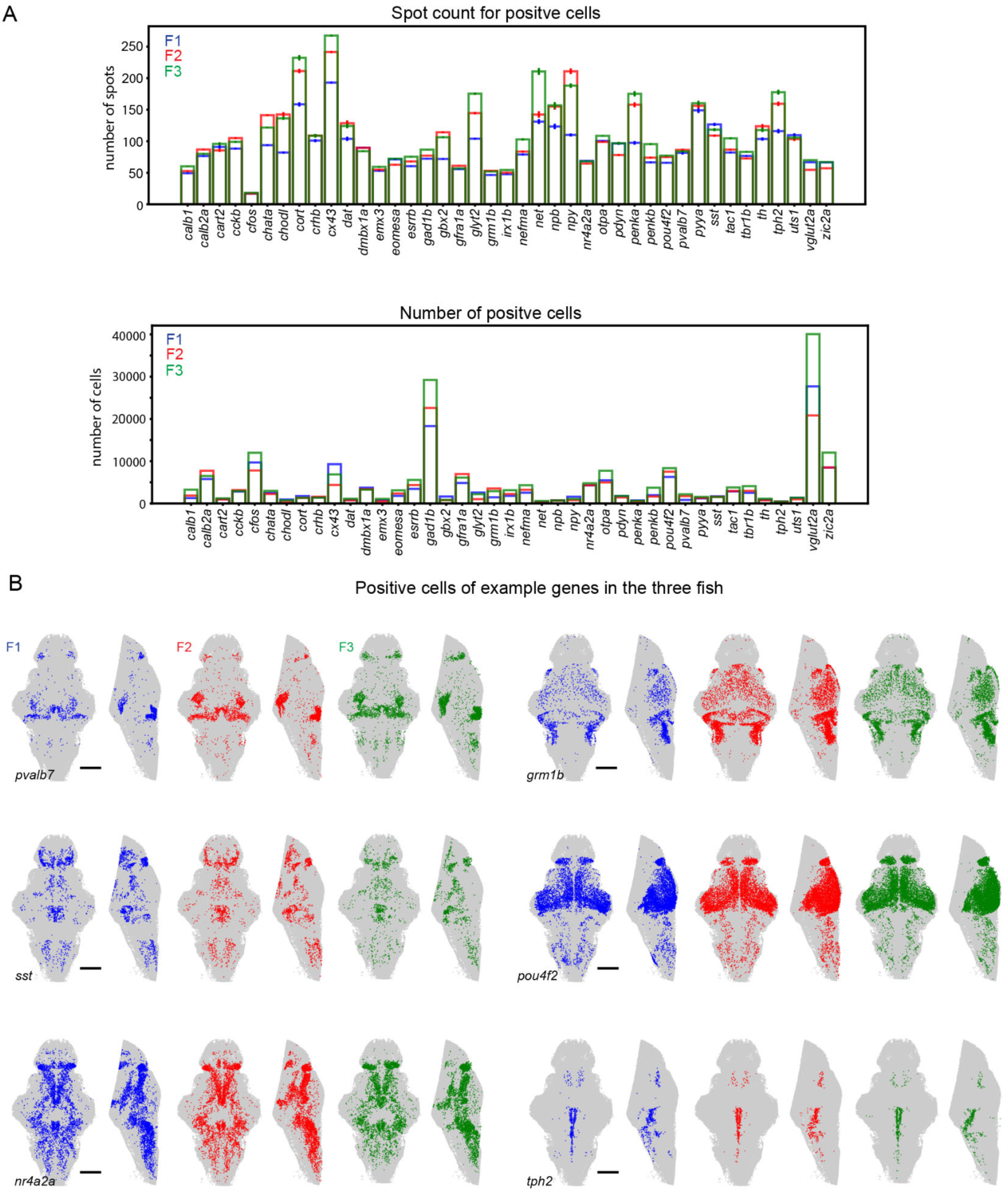
Gene expression distribution across samples. (A) Top: Spot count and positive cell number by animal. Left: Average spot count for the positive cells of each of the probed genes. Error bars indicate SEM. Bottom: Number of positive cells across the whole brain for each probed gene. Each color represents a single animal. (B) Distribution of positive cells across animals. Top and lateral view of the distribution of positive cells for 6 of the probed genes. Each color represents a single animal.

**Figure S5:**
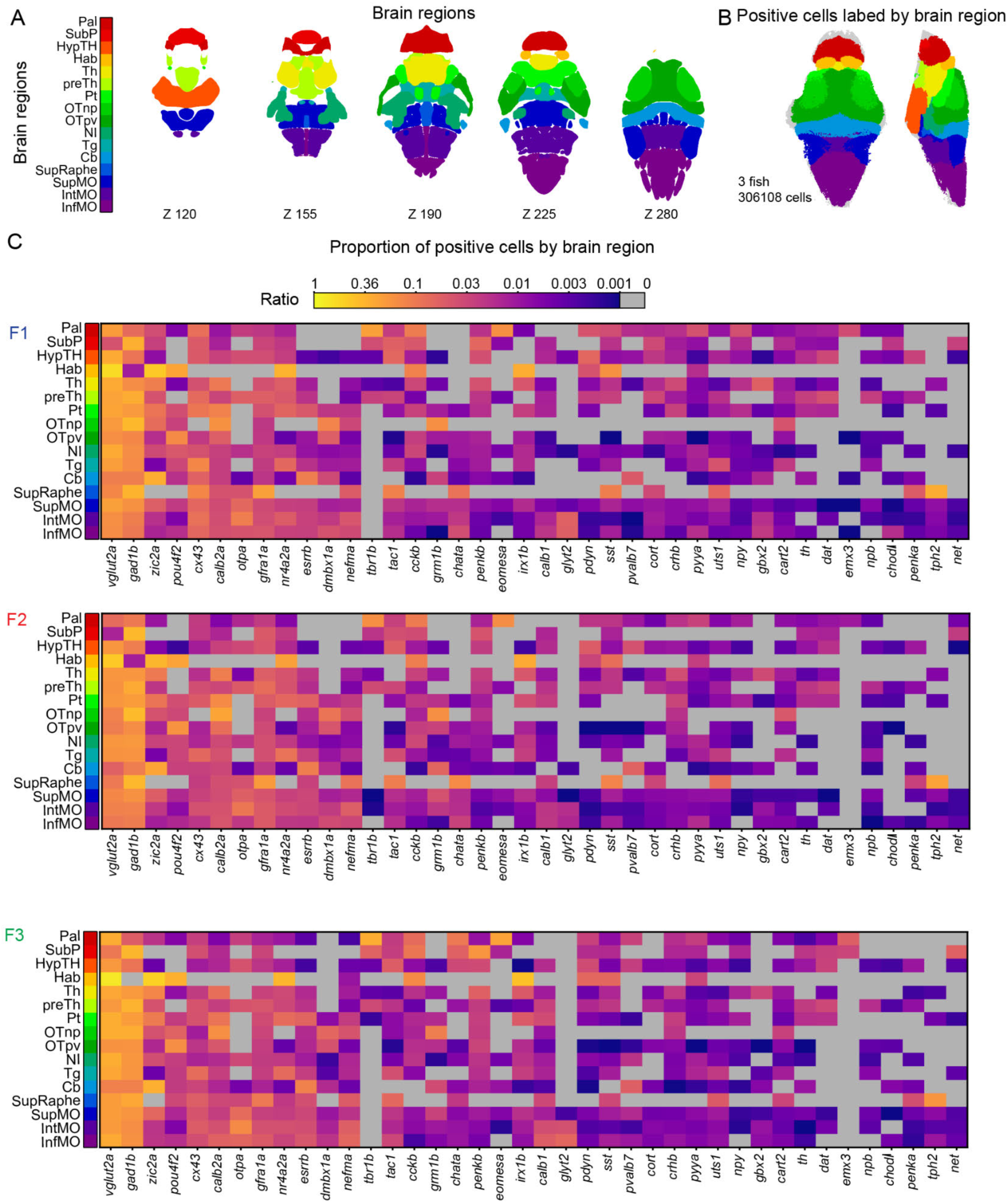
Gene expression distribution across brain regions. (A) Brain regions masks from the mapzebrain atlas, 5 different z planes are shown. (B) Positive cells and their location. Top and lateral view of the brain region localization of 306108 cells from 3 different animals that are positive for at least one of the probed genes. (C) Matrices with the average percentage of positive cells for the 40 genes (excluding *cfos*) within each of the 16 brain regions for each of the animals.

**Figure S6.**
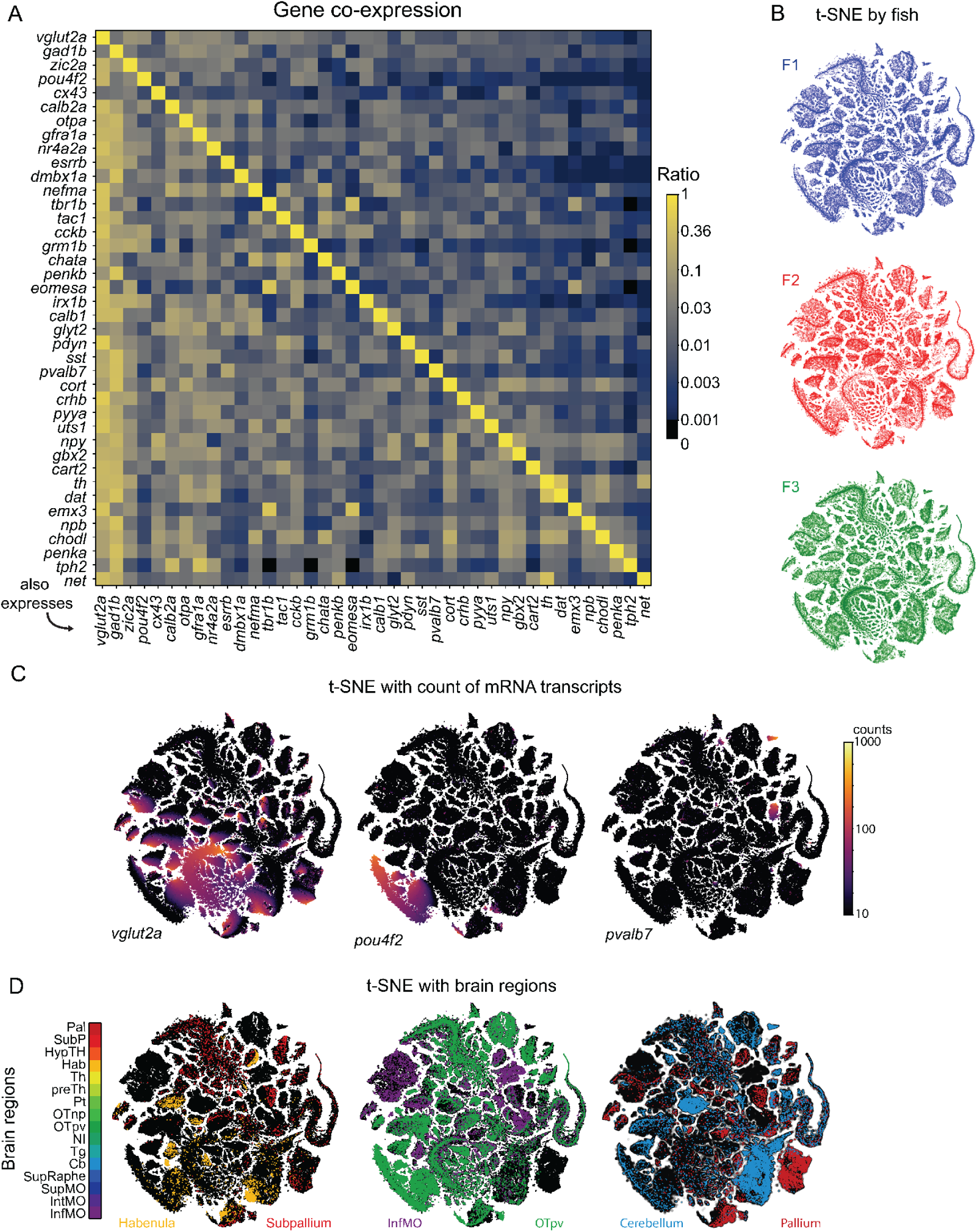
Co-expression and molecular subtypes (A) Matrix with the average proportion of cells that are positive for 2 genes across the 3 fish. Proportion of cells positive for genes in the two rows that are also positive for the genes in the columns. (B) Scatter plots with the cells of each of the animals in the T-SNE 2-dimensional embedding of the gene expression in Figure 4C. (C) Spot counts in the T-SNE 2-dimensional embedding of the gene expression in Figure 4C. The colormap indicates the spot count values for 3 example genes. (C) T-SNE 2-dimensional embedding of the gene expression in Figure 4C labeled with pairs of brain regions.

**Figure S7:**
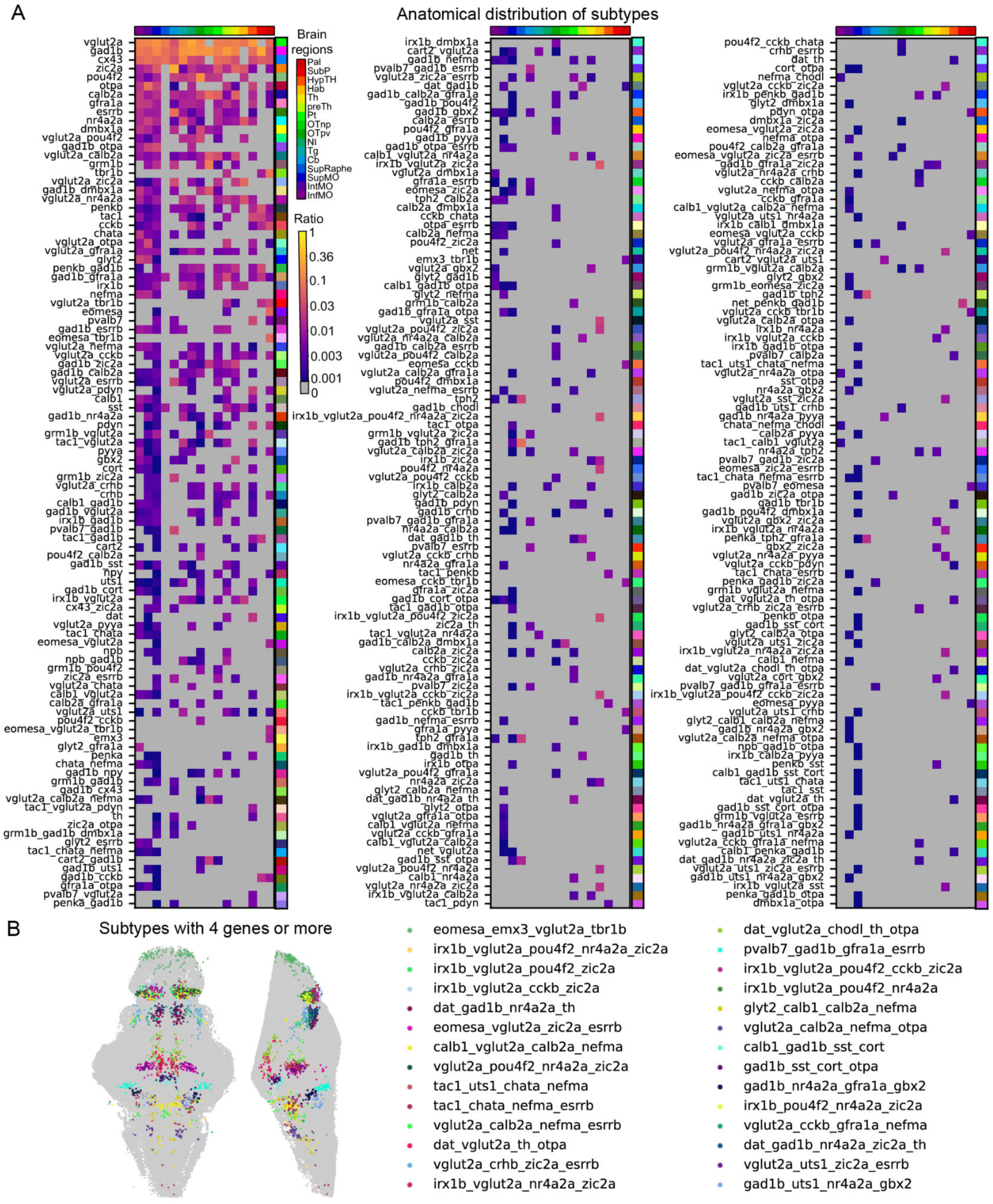
Co-expression and molecular subtypes (A) Matrix with the average proportion of cells that are part of the subtypes (rows) in each of the brain regions (columns). (B) Example subtypes with neurons that are positive for 4 or 5 genes.

**Figure S8:**
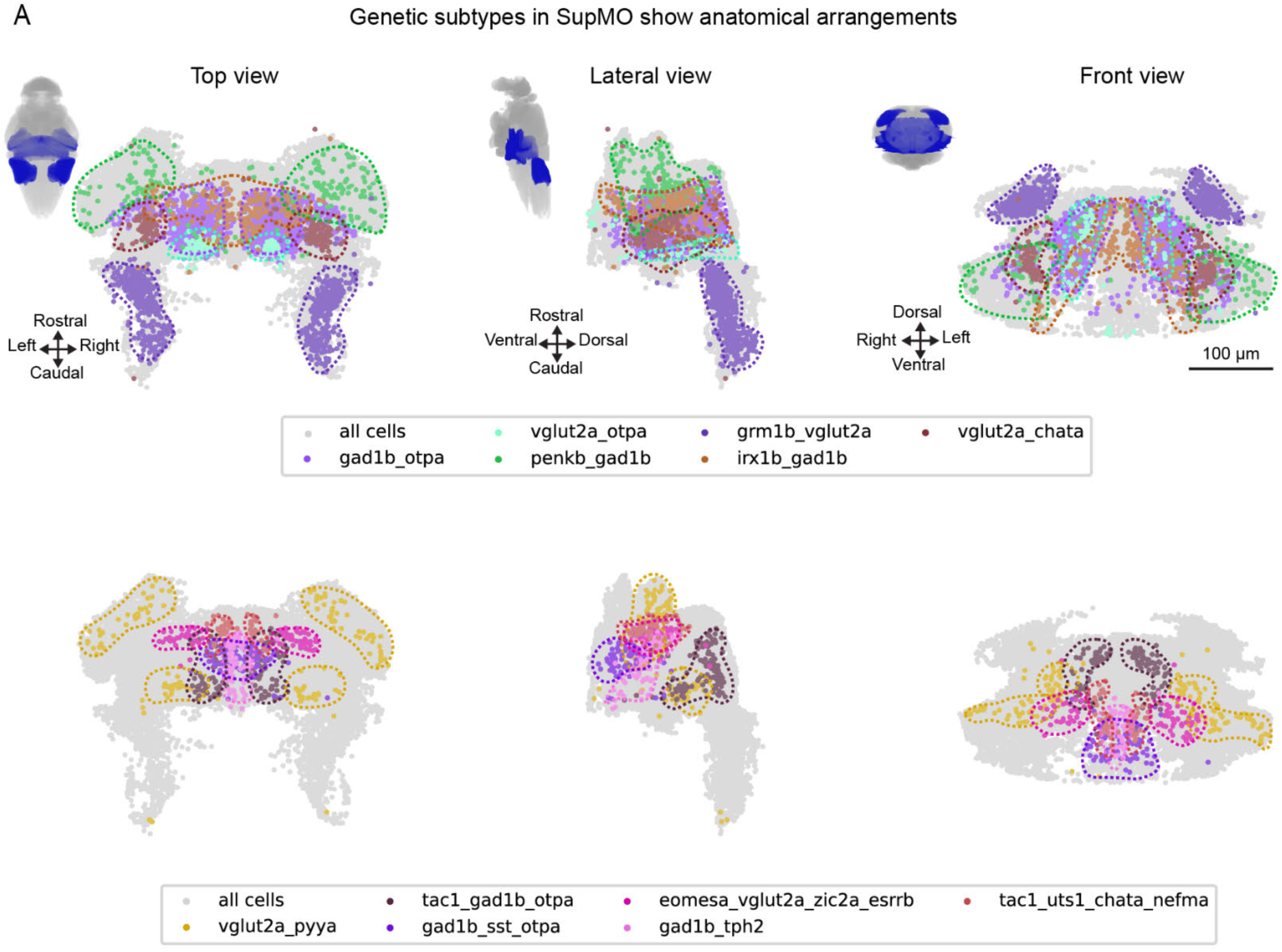
Molecular subtypes display different distributions that suggest anatomical organizations. (A) Twelve example molecular subtypes and the anatomical localization of their neurons seen from a top view (left), lateral view (middle) and front view (right). Some subtypes show a scattered and wide distribution (e.g. penkb_gad1b), while other show more compact organizations (e.g. vglut2a_chata, tac1_gad1b_otpa, tac1_uts1_chata_nefma). Some are organized in bilateral clusters (e.g. eomesa_vglut2a_zic2a_esrrb, vglut2a_pyya), in dorsoventral columns (e.g. gad1b_otpa, vglut2a_otpa, irx1b_gad1b) or in rostrocaudal orientations (grm1b_vglut2a). Some are in the midline, resulting in one compact population instead of 2 symmetrical ones in each hemisphere (e.g. gad1b_sst_otpa, gad1b_tph2).

**Figure S9:**
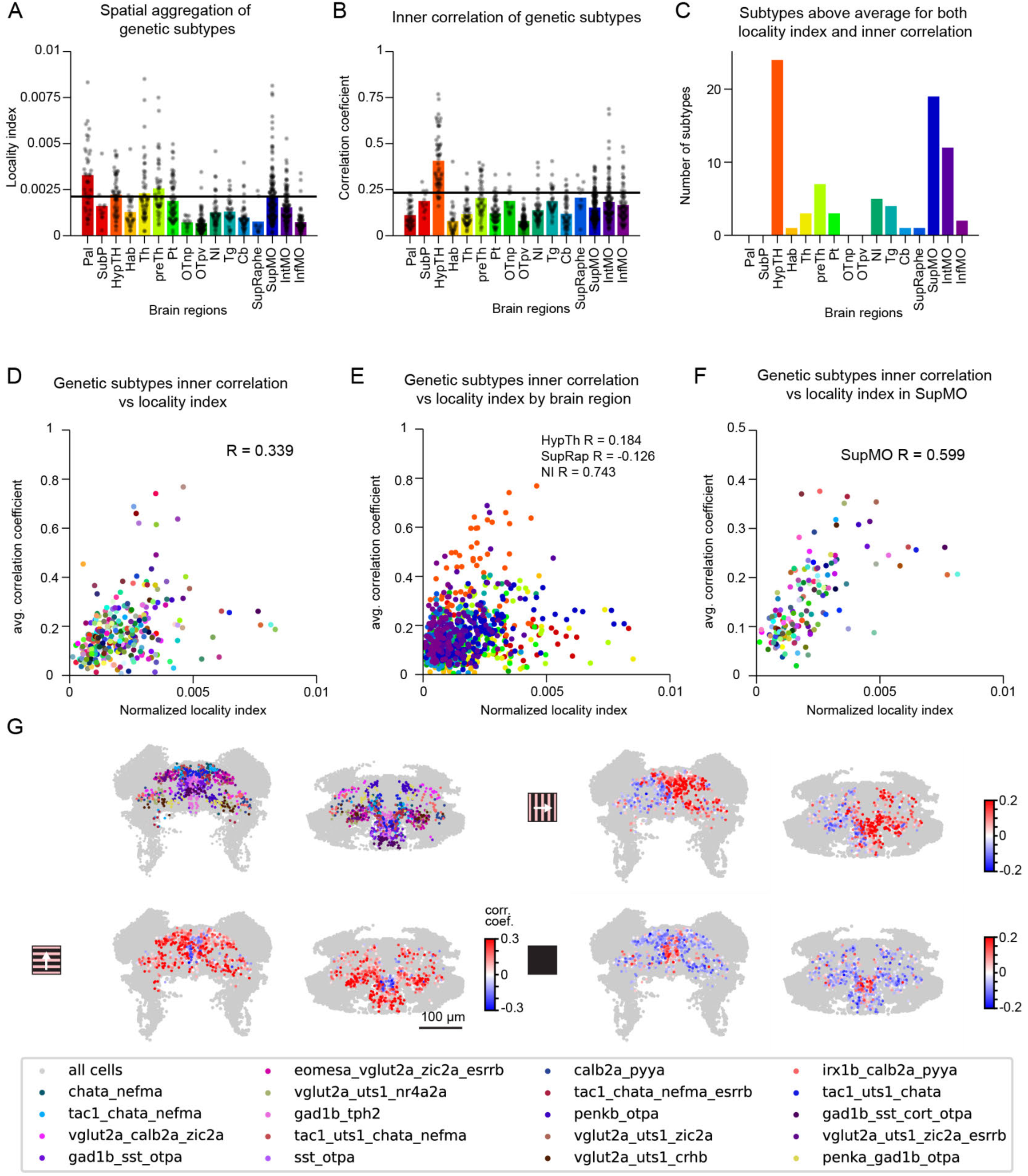
Compact subtypes are more coherent (A) Average locality index of the subtypes by brain region. The horizontal black line represents the brain-wide average. (B) Average inner correlation of the subtypes by brain region. The horizontal black line represents the brain-wide average. (C) Number of subtypes above average in both locality index and inner correlation by brain region. (D) Scatter plot showing the linear relationship (R=0.339) between the inner correlation of each subtype and their normalized locality index. For this analysis the average correlation and locality index are averaged across brain regions. This relationship suggests that more compact subtypes tend to have cells with more similar activity. (E) Scatter plots as in D, for each of the selected brain regions. (F) Scatter plot as in D, but for molecular subtypes found in the SupMO. (G) Top left: molecular subtypes in the SupMO which normalized locality index and inner correlation are above the mean (brain-wide). Bottom left: correlation of the activity to the forward visual motion presentation for the cells shown above. Top right: correlation of the activity to the rightward visual motion presentation for the cells shown in top left. Bottom right: correlation of the activity to the dark flash presentation for the cells shown in top left.

**Figure S10:**
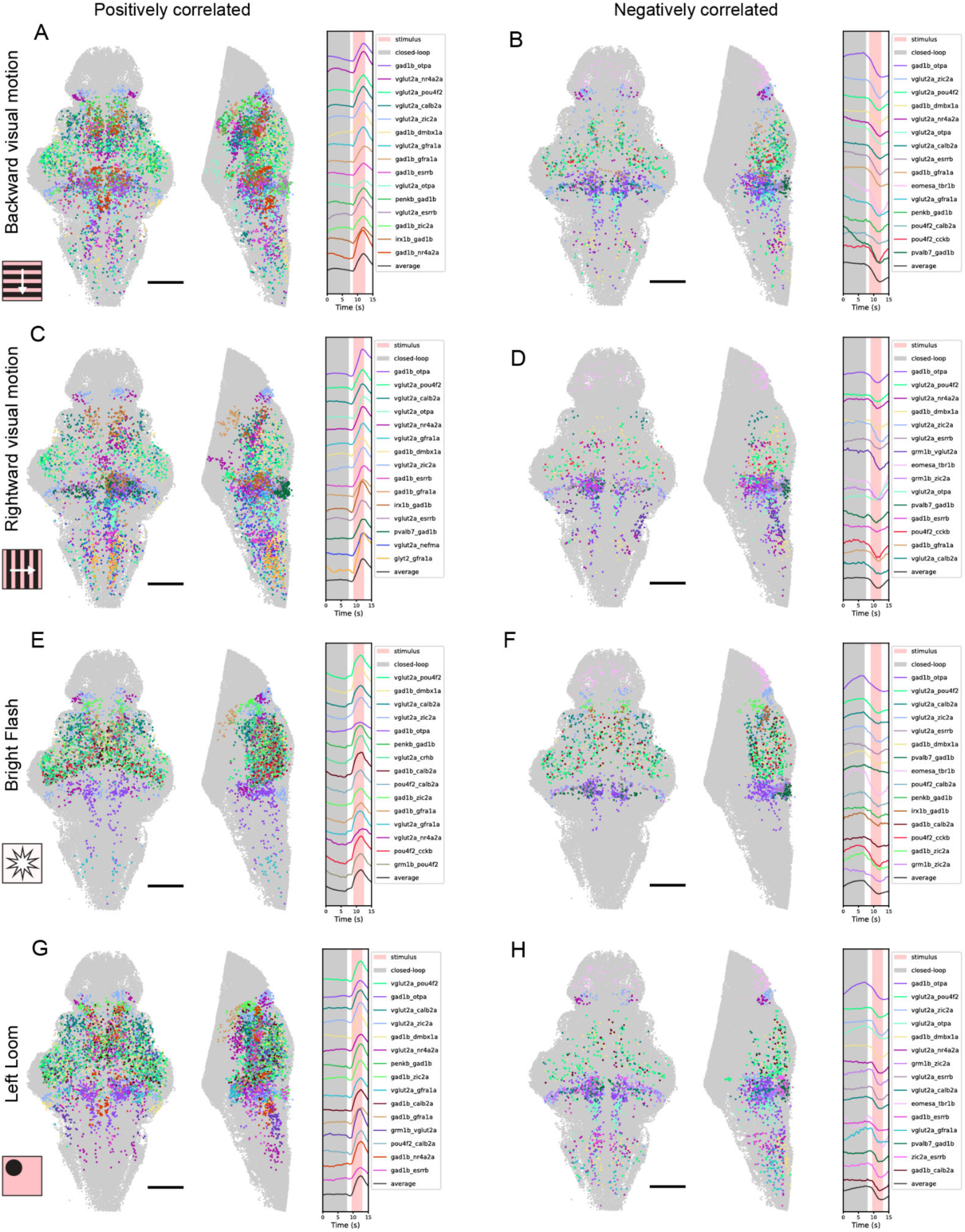
Anatomical distribution of example responsive subtypes (A), (C), (E) and (G). The 15 largest subtypes positive for more than one gene that are positively correlated with backward visual motion, rightward visual motion, bright flash and left loom. Left: Top view of the subtypes in their anatomical location. Middle: lateral view. Right: Average activity trace of each subtype. (B), (D), (F) and (H). The 15 largest multi-gene subtypes that are negatively correlated with backward visual motion, rightward visual motion, bright flash and left loom. Left: Top view of the subtypes in their anatomical location. Middle: lateral view. Right: Average activity trace of each subtype.

**Figure S11:**
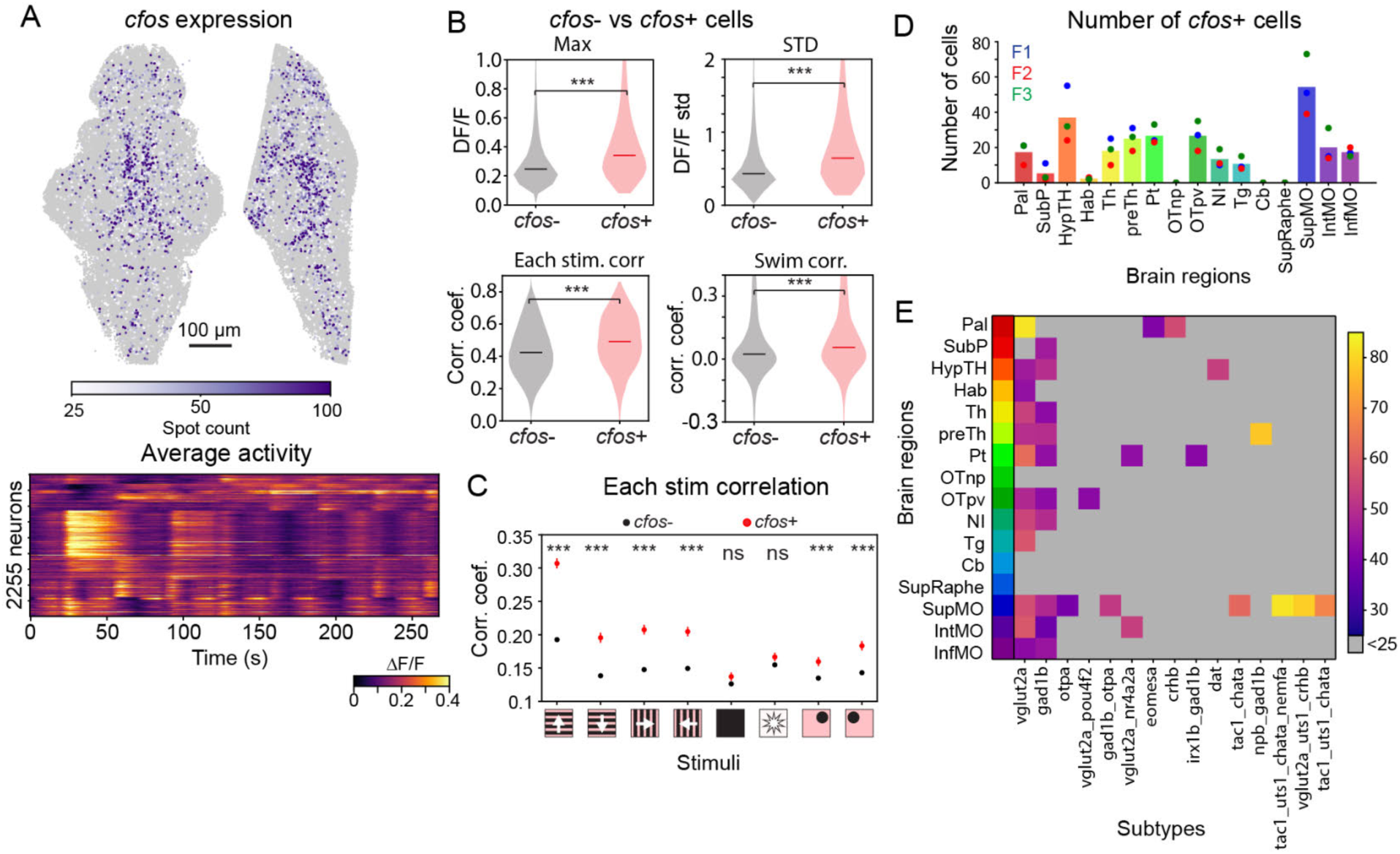
*cfos* expression in active cells (A) Top: Brain map with *cfos*+ cells that have a minimum of 25 spots. Bottom: trial average activity trace of each of the *cfos*+ neurons from the brain map above. Ordered by similarity in activity and binned every 10 neurons. (B) Differences between *cfos*- and *cfos*+ cells. Top left: Maximum ΔF/F; Top right: standard deviation of the ΔF/F. Bottom left: correlation coefficient values to stimulus presentations. Bottom right: correlation coefficient values to swim activity. (Mann-Whitney U test, ns p>0.05; * p<0.05; ** p<0.01; *** p<0.001). (C) Differences in correlation to stimuli. (Mann-Whitney U test, ns p>0.05; * p<0.05; ** p<0.01; *** p<0.001). (D) Average (across fish) number of *cfos*+ cells per brain region. (E) Matrix with the average number of *cfos* spots for each subtype within different brain regions. Each row represents one of the defined brain regions, each column represents one of the defined subtypes.

**Figure 12.**
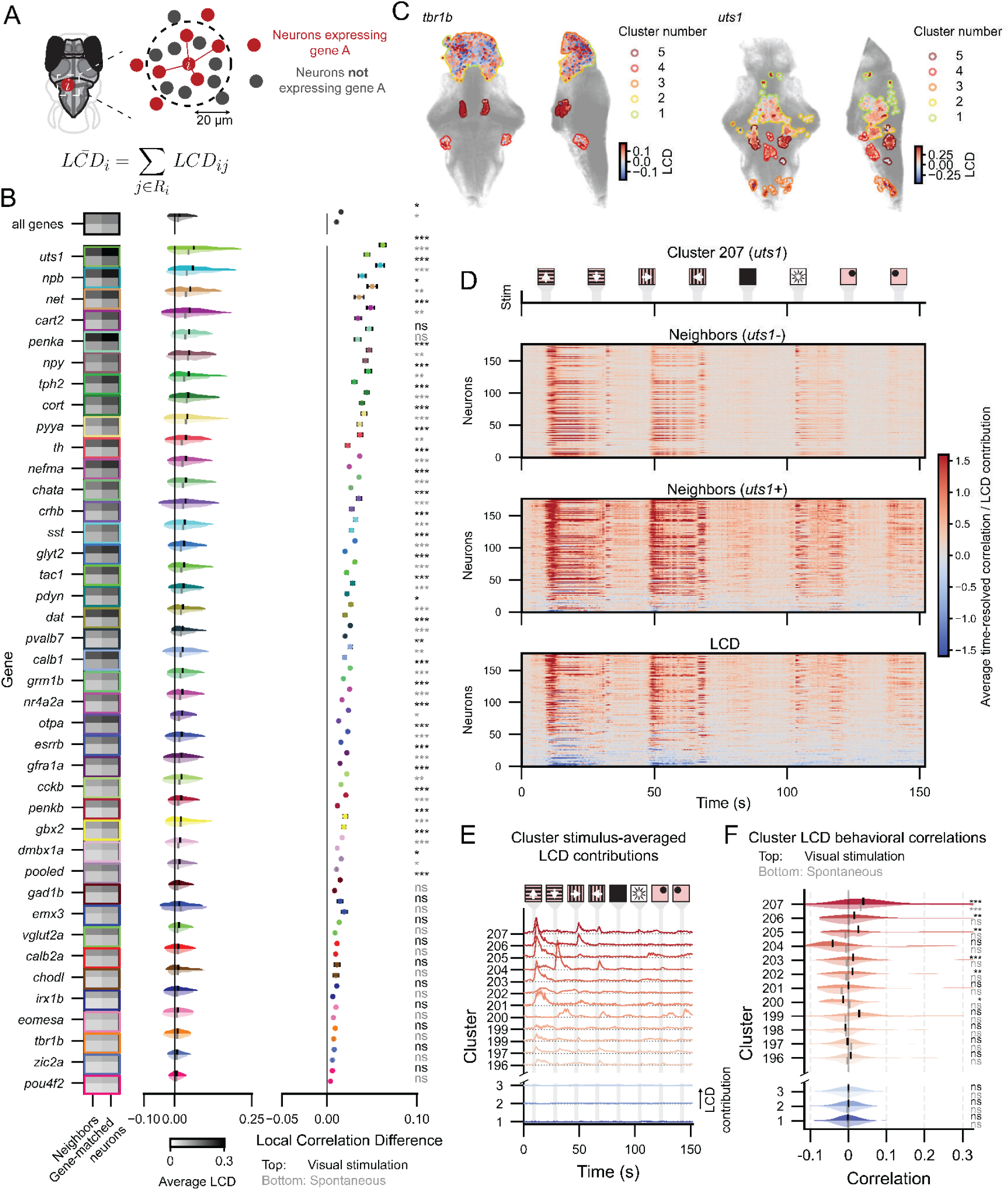
Local Correlation Differences (LCDs) A) Because individual pairwise LCD values can be noisy, we summarized LCD at the level of single neurons and spatial bins. For each neuron, we averaged *LCD_ij_* over gene-matched partners within a spherical neighborhood of radius 20um around that neuron, resulting in a neighborhood-averaged *LCD_i_* for each neuron expressing a given gene. B) LCD distributions for both the visual stimulation (top plots for each row) and spontaneous (bottom plots for each row) paradigms for all genes except *cx43* and *cfos* (extension of Figure 6C). First column: average correlation of neurons with neighbors of the same genetic type vs. remaining neurons; second column: LCD distributions for the same genes; third column: mean ± SEM and significance of these distributions. All columns are ordered according to the median LCD. (Two-sided Wilcoxon Signed Rank Test, Benjamini-Hochberg corrected across genes: ns p > 0.05; * p<0.05; ** p<0.01; *** p<0.001; Visual stimulation significances indicated in black, spontaneous in grey; across all genes, p=0.184 (visual stimulation), and p=0.222 (spontaneous); individual gene values in Data S1.) C) Spatial maps of LCD values for *tbr1b+* and *uts1+* cells across all fish, showing similar patterns across animals. Colors indicate the within-gene spatial clusters to which the neurons were assigned. D) Average contribution to the Pearson correlations from non-gene expressing neighbors (top), gene-expressing neighbors (middle), and their difference constituting the LCD contribution (bottom) for each time step for spatially localized cluster 207 (expressing *uts1)*. E) Stimulus-averaged LCD contributions per time step for the 12 clusters with the highest and the 3 clusters with the lowest median LCDs (colors matching Figs 6D-G). Shaded areas indicate ± SEM. F) Violin plots showing the distribution of correlations to the measured fictive behavior of the visually stimulated paradigm (top half of violins) and spontaneous paradigm (bottom half of violins) of the animal for the same clusters as shown in panel F. (Two-sided Wilcoxon signed-rank test, Bonferroni-corrected; per-cluster values in Data S1).

